# Motor neuron boutons remodel through membrane blebbing

**DOI:** 10.1101/2021.03.07.434250

**Authors:** Andreia R. Fernandes, César S. Mendes, Edgar R. Gomes, Rita O. Teodoro

## Abstract

Wired neurons form new presynaptic boutons in response to increased synaptic activity, but the mechanism by which this occurs remains uncertain. The neuromuscular junction (NMJ) is a synapse formed between motor neurons (MNs) and skeletal muscle fibers and is critical for control of muscle contraction. Because *Drosophila* MNs have clearly discernible boutons that display robust structural plasticity, it is the ideal system in which to study bouton genesis. Here we show using *ex-vivo* and by live imaging that in response to depolarization, MNs form new boutons by membrane blebbing, a pressure-driven mechanism used in 3-D migration, but never described as a neuronal remodeling strategy. In accordance, F-actin is decreased during bouton growth (a hallmark of blebs) and we show that non-muscle myosin-II (a master regulator of blebbing) is recruited to newly formed boutons. Furthermore, we discovered that muscle contraction plays a mechanical role in activity-dependent plasticity, promoting bouton addition by increasing MNs confinement. Overall, we provide a novel mechanism by which established circuits create new boutons allowing their structural expansion and plasticity, using trans-synaptic physical forces as the main driving force. Understanding MN-muscle interplay during activity-dependent plasticity can help clarify the mechanisms leading to MN degeneracy observed in neuromuscular diseases.

---

The establishment of synapses enables the wiring and communication of neurons into circuits, that when activated allow perception, learning and memory, and locomotion. These circuits are determined by genetic programs during development^1^, but can be modified throughout adulthood by activity-dependent plasticity mechanisms to support information storage and environmental adaptation^2–4^. Presynaptic boutons are conserved round specializations of axonal membrane where neurotransmission takes place^5–7^ and their presence is a good indicator of the existence of a functional synapse^8,9^. Boutons are highly plastic structures that undergo rearrangements in response to changes in neuronal activity, including synapse formation and elimination. By contributing with significant and stable rewiring of circuits, boutons can act as powerful modulators of neuronal output^10–13^. Further supporting their importance to maintain network function, several studies highlight a link between presynaptic-malfunction and neurodegenerative disease etiology^14–16^.

Despite bouton’s importance, there is a lack of understanding about the mechanism of their formation in wired neurons. During embryogenesis, neuronal axons extend towards their synaptic targets through highly dynamic structures comprised of filopodia and lamellipodia, called growth cones (GC), which upon arrival to their destination differentiate into round boutons creating highly specific connections^17,18^. After this period, neurons are still able to remodel their structure as the animal grows, adding new boutons in response to developmental cues and to acute external stimuli^19–23^. However, the mechanism by which boutons form in these later stages remains elusive.

The *Drosophila melanogaster* larval NMJ has been widely used as an *in vivo* model to discover the machinery that regulates bouton formation^24,25^. Other than developmental bouton addition, which occurs throughout the larval stages, the advent of simple stimulation paradigms applied to dissected larval preparations, or intact larvae, created an opportunity to study in more detail the mechanisms required for activity-dependent bouton formation. Furthermore, combining these protocols with live imaging studies showed that activity-induced presynaptic morphogenesis may be different from the embryonic GC^26,27^.

It has been shown that new boutons form in a few minutes mostly while the muscle is contracting^27^, with no indication of structures similar to filopodia or GCs giving rise to new boutons^26^. Instead it was observed that new boutons emerge from the presynaptic arbor by budding off round varicosities of membrane^27^. These studies also revealed that, in addition to microtubule remodeling^28–34^, new bouton growth induced by elevated activity is associated with actin cytoskeleton dynamics^27,35–37^ and requires proteins that regulate synaptic vesicle-machinery^38–40^ suggesting that neuronal remodeling is likely coupled to synaptic transmission. Additionally, several signaling pathways have been shown to be implicated in the causal relationship between synaptic activity and bouton formation suggesting that activity-dependent plasticity at the NMJ results from a complex sequence of factors that are involved in the communication between MNs, muscle and glial cells^25^. Still, the molecular mechanisms and cytoskeletal dynamics used by MNs to coordinate their activity with its cellular partners, to initiate structural plasticity, are poorly understood. Recent studies show that synaptic changes prompted by activity suggest dynamic cell interactions involving cell surface and extracellular matrix adhesive proteins^36,41–49^. Because, coupled to the cytoskeleton, these molecular players are known to govern trans-synaptic forces, it has been proposed that synapse formation can be mechanochemically regulated^50,51^. Yet, the role of tissue mechanics for synapse formation and plasticity is largely unexplored.

Here, we report for the first time that neuronal remodeling at the *Drosophila* NMJ occurs by membrane blebbing, a pressure-driven mechanism used in 3-D migration, and that this process requires an interplay of mechanical forces between neurons and muscle to coordinate activity-dependent bouton formation *in vivo*.

## Results

### Activity-dependent bouton formation mechanistically resembles blebbing migration

To study bouton formation in the context of presynaptic remodeling, we used the *Drosophila* third instar larval NMJ, where developmental bouton addition is nearly complete, and fast formation of new boutons can be prompted by patterned High-K^+^ stimulation^26,38^. Our strategy was to follow boutons formed in response to elevated neuronal activity by live imaging with high-temporal resolution (in secs) in larvae subjected to acute stimulation of structural plasticity (**Supplementary Fig. 1a**). Based on an earlier study that reported bouton formation while the larvae was contracting^27^, we decided to image the NMJs of unasthetized larvae hypothesizing that muscle activity could contribute to this process. We imaged third instar larvae NMJs expressing CD4-tomato (CD4-Tom) in all neurons using NSyb-Gal4 to label the presynaptic membrane **(Fig. 1a and Movie 1 and 2).** Video analysis showed that the membrane dynamics that accompany activity-dependent bouton addition are clearly distinct from the embryonic GC, as described^27^. We identified two main types of bouton appearance (**Fig.1a**), mainly forming from preexisting boutons: first, with rapid membrane expansion with bursting growth that was frequently associated with clear muscle contraction events (**Movie 1**); and second, with gradual membrane flow and slow elongation associated with subtle muscle movements (**Movie 2**). Although we do not exclude more types of bouton formation, our data indicates that activity-induced boutons do not emerge from lamellipodia and/or filopodia precursors. Instead, we observe that boutons always emerged as large spherical expansions of MN membrane and often their growth was a rapid event, ranging from less than 1 min to a few mins. The median time for bouton formation observed with CD4-Tom was 5min32s (**Fig. 1b**). On several occasions, a bright membrane punctum was visible at the base of newly formed boutons, suggestive of local intracellular vesicle dynamics.

**Fig.1:**
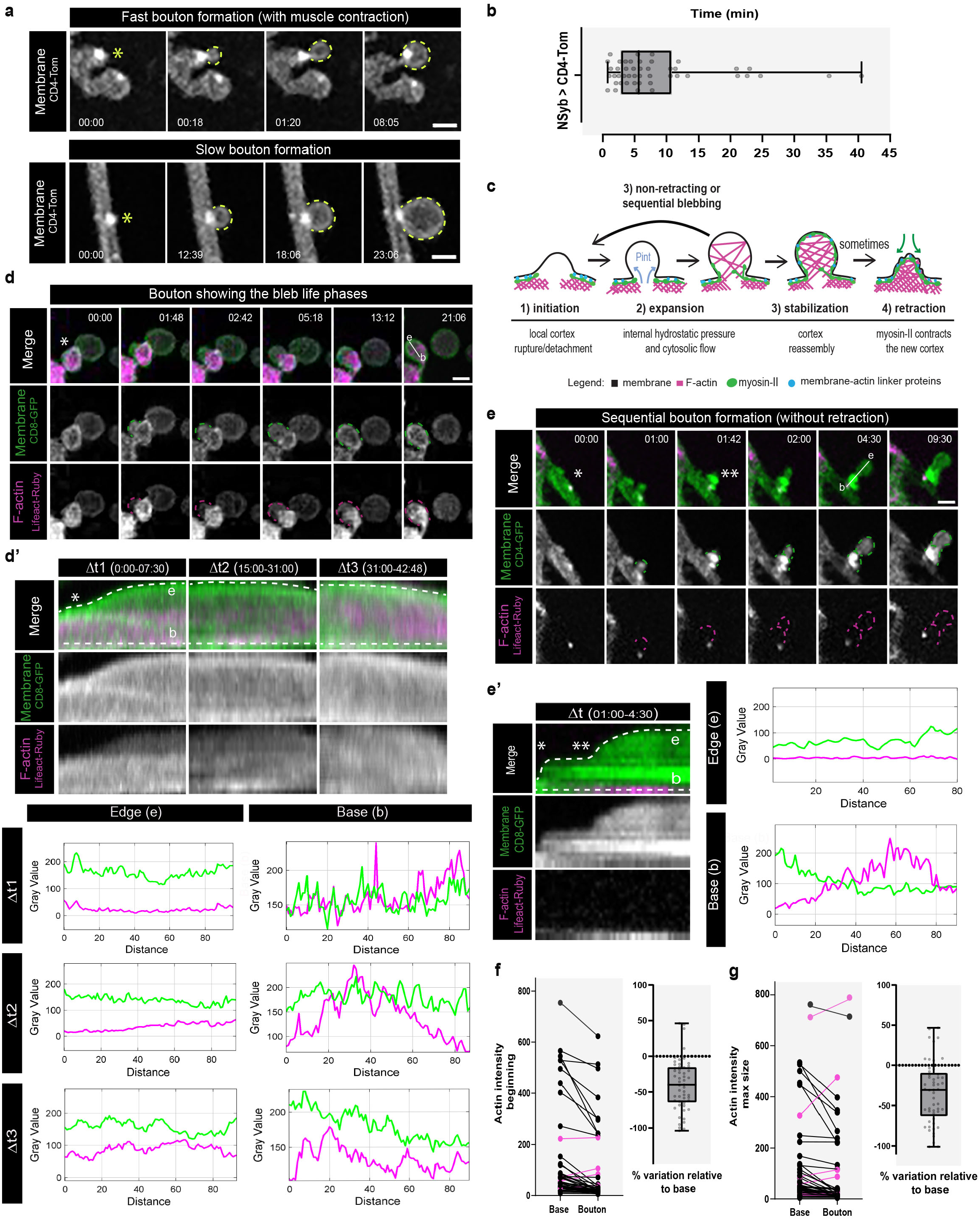
Activity-dependent bouton formation mechanistically resembles blebbing migration. **a,** Time-lapse images of bouton formation. Examples of fast bouton formation (top), frequently observed with muscle contraction, and slow bouton formation (bottom). Neuronal membrane was labed with UAS-CD4-Tom under the control of NSyb-Gal4 (pan-neuronal driver). Scale bars, 2 μm. Asterisk indicates where bouton will emerge, **b,** Boxplot (min to max) with times of bouton formation across experiments. Median is 5min38s. All data points are represented. n=13 larvae, 46 boutons, **c,** Schematics of bleb life cycle. Non-motile cells showing: 1) initiation; 2) rapid expansion (secs); 3) stabilization; 4) slow retraction (a few min). Retraction is rarely seen in motile cells, **d,e,** Time-lapse images of new boutons showing the bleb life phases. Neuronal membrane and F-actin were labeled with UAS-CD8-GFP (or CD4GFP) with UAS-Lifeact-Ruby, both under the control of NSyb-Gal4. Scale bars, 2 μm. Asterisk indicates where bouton will emerge, **d,** Example of bouton formation and subsequent stabilization, **e,** Example of sequential bouton formation without retraction, **d’,e’**, Kymographs from d and e. kymographs for distinct Δt’s to highlight the dynamic nature of actin during bouton growth. Dotted lines along the bouton edge (e) and base (b) and the correspondent intensity values for membrane and F-actin below, d’ Initially actin is scarce (Δt1) but is subsequently assembled inside boutons and accumulates when bouton size stabilizes (Δt2 and Δt3). e’, Actin is absent in new boutons and an F-actin punctum persists at the base, **f,g,** Plots with absolute values for F-actin intensity (left) measured in boutons and correspondent bases at beginning (**f**) or max size (**g**), confirm low actin in new boutons. Boxplot (min to max) showing % of variation in F-actin intensity (right) in each bouton compared to its base (used as 100% reference). Line represents the median. All data points are represented. n=14 larvae, 53 boutons.

Given the morphological resemblance with yeast budding, a budding mechanism has been previously proposed to explain activity-dependent bouton formation at the *Drosophila* NMJ^52^. However, budding is mostly a slow process taking hours to occur and can only be as short as 1,5h under optimal conditions^53^. Our imaging data suggests that bouton formation is a more dynamic process that strongly resembles cellular blebbing. Blebs are round membrane protrusions extruded by intracellular hydrostatic pressure that some cells use, in combination or in alternative to lamellipodia, to migrate in 3-D environments^54,55^, but have never been reported to be used in neurons as a mechanism for structural plasticity. Membrane blebbing is evolutionarily conserved and is favored by conditions of high confinement and contractility^54^. Blebbing is in a large extent, a physical, rather than a biochemical process, and it occurs over time- and length-scales that have been little investigated in animal cells and *in vivo*^56^. Nevertheless, a characteristic feature of blebs is the rapid change in cell shape in the site where the protrusion occurs, which can occur in around 30s^56^. To assure that the transmembrane tag used, CD4-Tom, did not interfere with bouton formation, we compared this genotype with wild-type (*w^1118^*) and neuronal expression of cytosolic LexAop-mCherry under the control of DVGlut-LexA. Bouton frequency after stimulation was identical in the tested genotypes (**Supplementary Figs. 1 b-d**). Interestingly, the time of bouton formation was slightly increased in neurons expressing CD4-Tom (**Supplementary Figs. 1 e,f**) in the membrane, compared to neurons that expressed cytosolic mCherry, suggesting that CD4-Tom, which is an external tag, may delay bouton growth, possibly by interacting with the extracellular matrix. Since we may be overestimating, it is possible that boutons can form even faster (**Fig.1b**). We found no differences in new bouton area when using an external versus internal tag to label neuronal membrane (**Supplementary Figs. 1 g,h).**

Since rapid dynamics and morphological features were reminiscent of membrane blebbing, we sought to test if *Drosophila* MNs adapted this mechanism to remodel in conditions of intense neuronal activity. Non-motile cells blebs undergo fast expansion-retraction cycles (**Fig. 1c**), while migrating cells rarely retract and usually form persistent and even sequential blebs that eventually become stabilized^57^. Either way, a big difference between blebs and other membrane protrusions, such as lamelipodia or filopodia, is that bleb growth does not rely on actin polymerization but is driven primarily by a flow of cytosolic fluid across a local weakening (rupture or detachment) in the cell actomyosin cortex^54^. As a result, the main hallmark of blebs is that they are initially depleted of an actin cytoskeleton (beyond the fluorescence detection limit), but blebs can subsequently assemble the cytoskeleton to stabilize or retract^56,58^. Accordingly, it was shown that the actin cytoskeleton can undergo significant reorganization during a bleb’s lifetime^57^, which lasts only a few mins^54^. We used animals expressing CD4-GFP and Lifeact-Ruby in neurons to follow both membrane and F-actin dynamics in real-time after High-K^+^ stimulation (**Fig. 1d-e’** and **Movies 3-5**). This experiment showed that new boutons have the hallmark of blebs: membrane growth without prominent F-actin. Furthermore, we could observe bouton formation with and without retraction, identical to the bleb types previously described (**Figs. 1d-e’** and **Supplementary Figs. 2a,a’; Supplementary movies 3-5**). We quantified actin content in the new boutons, in the beginning and at maximum size, by establishing the difference between the base and the edge of the bouton, and showed that in growing boutons actin content was consistently found to be lower than in base (92% boutons at beginning, median −40%; 85% boutons at maximum size, median −31%) (**Figs. 1f, g** and **Supplementary Fig. 2f**). In several boutons we could detect reductions in actin that were superior to 50%, compared to basal actin. On the other hand, in the modest number of boutons that formed with actin higher than base (up to 15% boutons), these gains in actin were inferior to 50% compared to basal actin, suggesting that even in these cases low amount of actin was recruited as these boutons formed. We analyzed several aspects of bouton formation dynamics and its relationship with actin (**Supplementary Figs. 2 and 3**), overall concluding that in the majority of cases, actin is very low when boutons first form, remaining low throughout the process, even though we did observe several instances of actin repolymerization of the cortex (**Supplementary Figs. 2b and 3c**). The time for bouton formation seemed independent of the actin content, although there seemed to be a non-statistical trend towards boutons with increasingly higher actin content showing increasingly higher times to form (**Supplementary Figs. 2c and 3f**). Altogether, our data suggests that actin polymerization is not the main driving force for bouton growth. Still, we observed F-actin puncta at the base of new boutons in 26% of events **(Fig. 1e** and **Supplementary Fig. 3j),** which were possibly related with actin flow into boutons **(Supplementary Figs. 3k,l).** Interestingly, actin puncta occurred preferentially in cases where threads of boutons formed rather than with single events **(Supplementary Figs. 3m-o).** This finding was consistent with previous report that spots of GFP-moesin, an actin binding protein that localizes to cortical F-actin, were associated with sites of bouton growth^27^. Additionally, it suggests that local actin rearrangements may occur to sustain rapid and successive bouton formation as it happens in blebbing cells.

To support this live data, we analyzed fixed larvae expressing Lifeact-GFP under the control of NSyb-Gal4, confirming that new boutons lack actin (**Supplementary Fig. 4a).** Interestingly, we also found that new boutons are depleted of filamin (**Supplementary Fig. 4b**), an F-actin crosslinking protein that links the membrane and intracellular proteins to actin. Since filamin in cells is generally distributed throughout large actin-base structures, such as lamellipodia and cortical actin^59,60^, together with our movies this finding supports that new boutons form with a weaker actin structure, identical to membrane blebs.

### Manipulation of actin dynamics during acute stimulation regulates bouton size

As actin-machinery plays a critical role in blebbing, and because our live imaging indicates that actin is dynamic throughout bouton growth, we wanted to pharmacologically assay if moderately shifting actin to a less or more polymerized state (**Fig. 2a**) changed activity-dependent bouton formation, and, if so, in which bleb phase. We used the number of F-actin puncta observed in boutons attached to the main axonal arbor as readout for the effect of the drugs on actin dynamics. Application of 10 μM Latrunculin B (LatB) to the stimulation solutions rapidly dispersed actin puncta (**Fig. 2a, top**), suggesting that the rate of depolymerization is increased. Whereas Jasplakinolide (JAS) treatment, stabilized existing actin puncta and caused the formation of new puncta indicating that it promoted actin polymerization (**Fig. 2a, bottom**). Contrary to what was previously observed for Latrunculin A^27^, a more potent inhibitor of actin polymerization, treatment of MNs with Latrunculin B did not change bouton frequency after High-K^+^ depolarization compared to the control conditions (HL3.1 only and DMSO vehicle) (**Figs. 2 b,i and Supplementary Fig. 5a**), however new bouton size was significantly increased (**Figs. 2 b-d**). This result suggests that a mild disruption of actin polymerization is not sufficient to impair acute structural plasticity, possibly because actin presence is not essential during the initial stages of bouton formation. Likewise, it was shown that local application of Latrunculin B does not change bleb frequency in cells but leads to a global increase in the bleb size^61^, in agreement with our data. Moreover, this study also reported that bleb dynamics ceased (slower rate) in the treated area, resuming only after removal of Latrunculin B, implying that actin polymerization is required for bleb retraction^61^.

**Fig.2:**
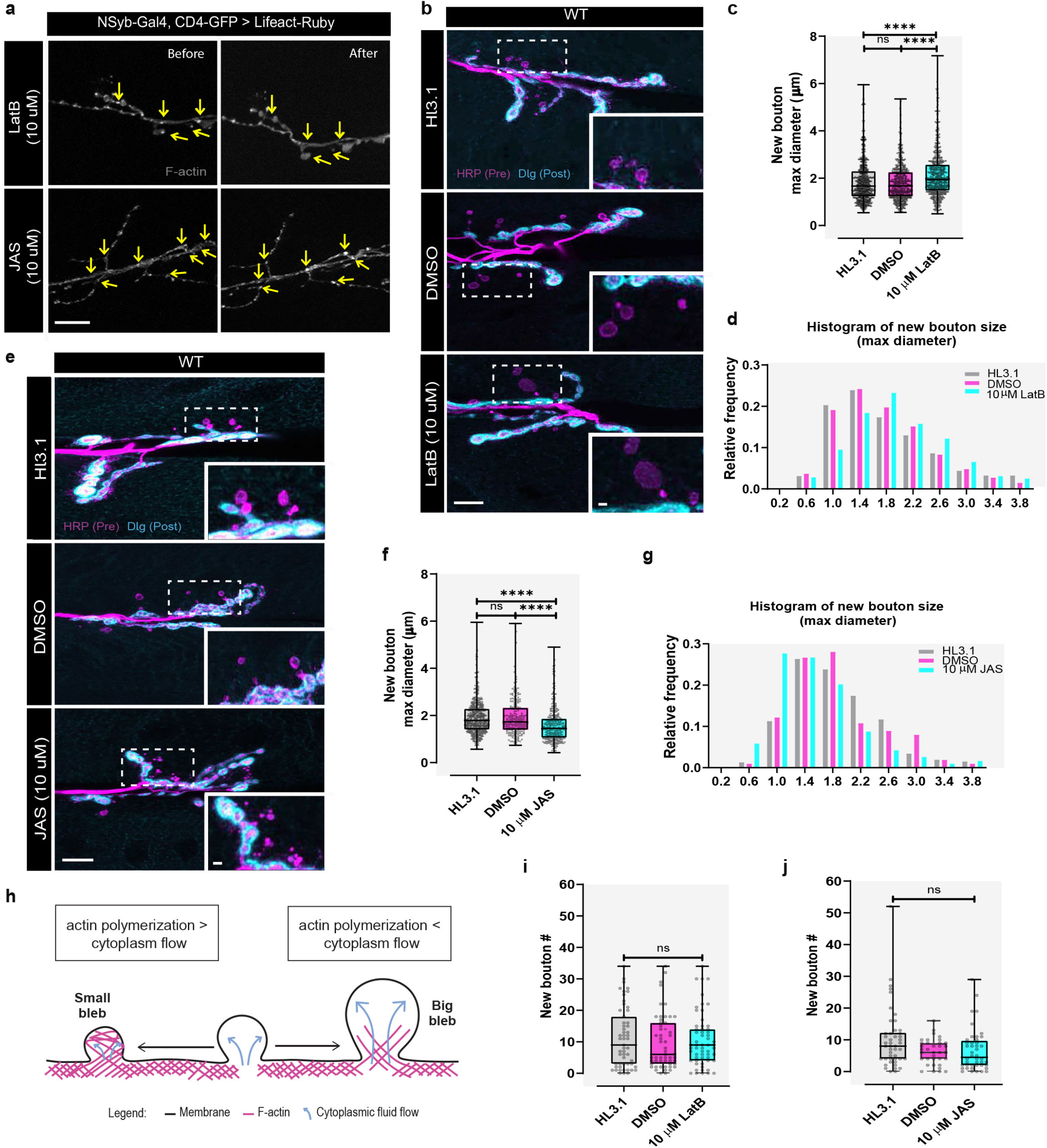
Manipulation of actin dynamics during acute stimulation regulates bouton size. **a,** Treatment of larvae with Latrunculin B (LatB, top) or Jasplakinolide (JAS, bottom). Representative images before and after drug application (15 min). LatB application lead to the rapid dispersion of F-actin puncta. After JAS treatment, F-actin puncta persisted/be- came brighter and new puncta appeared in some regions. F-actin was visualized by expression of UAS-Lifeact-Ruby under NSyb-Gal4 and F-actin puncta in boutons attached to the main arbor were used as readout of drugs effect at the NMJ. Scale bar, 10 μm. Yellow arrows indicate altered regions after incubation with the drug, n >6 larvae per condition, **b**, Images of WT animals (*w1118*) treated with HL3.1 alone, DMSO (solvent control) or 10 μM LatB after High-K+ stimulation. Scale bar, 10 μm. **c**, Boxplot (min to max) showing new bouton size (max diameter) is increased after LatB treatment. Line is median, d, Histogram showing relative frequency of new bouton sizes after LatB application, **c,d** n > 8 larvae ≥ 55 NMJs, ≥ 629 boutons for each condition and 3 biologically independent experiments, **e,** Images of WT (W1118) animals treated with HL3.1 alone, DMSO (solvent control) or 10 μM JAS after High-K+ stimulation, **f,** Boxplot (min to max) new bouton size (max diameter) is decreased after LatB treatment. Line is at median, g, Histogram showing relative frequency of new bouton sizes after JAS application, **f, g**, n≥ 6 larvae, ≥ 35 NMJs, >214 boutons for each condition and 3 biologically independent experiments, h, Schematics of bleb size regulation. Bleb size is controlled by a balance between cytoplasm flow driving expansion and actin polymerization that stops growth, **i** and **j,** Boxplot (min to max) showing that new bouton number is not altered either by LatB (**i**) or JAS (**j**). Line is at median, n > 8 larvae >55 NMJs (LatB) or n > 6 larvae, ≥ 35 NMJs (JAS). Presynaptic membrane labeled with HRP (magenta) and postsynaptic membrane labeled with Dig (cyan). New boutons (asterisk) were identified by the lack of Dig. Zoom of new bouton examples (corresponds to dashed square). Scale bar, 2 μm. Statistical significance was determined with non-parametric Kruskal-Wallis test (two tailed); ****p<0.0001, ns is non-significant.

On the other hand, treating MNs with JAS, leads to more variability in the response. Although we could observe that several NMJs produced fewer boutons, akin to what has been reported for JAS effects at the NMJ^27^, we also observed that other NMJs were unaffected by JAS treatment. Altogether, across experiments, JAS did not change bouton frequency after stimulation (**Figs. 2 e,j** and **Supplementary Fig. 5b**). Yet, we observed a remarkable phenotype in these NMJs, which was the presence of very small new boutons, usually clustered together in discrete regions (**Fig. 2e**). Additionally, we found new bouton size was significantly smaller after JAS treatment (**Figs. 2 f,g**). Altogether, our results suggest that bleb size seems to be determined by a balance between the initial growth rate (the rate of volume expansion due to cytoplasm flow) and the time needed for actin to repolymerize at the bleb membrane, which can be changed according with prevailing local actin dynamics. While JAS has been reported to either cause or block blebbing^62,63^, one hypothesis is that in cases where the flow rate of cytoplasm is insufficient to outpace local actin polymerization it could result in small or no blebs, whereas cases where actin filaments are highly stabilized would also restrict bleb formation. Supporting this hypothesis smaller boutons appear frequently together or in nearby regions, suggesting that they were formed during the same event, sequentially.

Altogether, we showed that new boutons show the actin cytoskeletal hallmark and dynamics (**Figs. 1 and 2**) compatible with being *bona fide* blebs, which supports the hypothesis that MNs may have adopted blebbing to rapidly modulate bouton formation in response to increased activity.

### Myosin-II, a master regulator of blebbing, is dynamically recruited to new boutons

Our data shows that activity-dependent bouton formation mechanistically resembles blebbing migration in which cells need to dynamically change their form – creating blebs – to move forward or invade tissues. Although blebs can be triggered in different ways, their generation is favored in conditions of high cortical tension, which is primarily generated through non-muscle myosin-II (myosin-II) activation. Indeed, drugs that relax the cortex by strongly constraining actin or myosin-II impair blebbing, suggesting that membrane detachment and bleb inflation are driven by intracellular pressure transients generated by contractions of the actomyosin cortex^58^ (**Supplementary Fig. 6a**). To study myosin-II distribution at the NMJ, under basal conditions and post High-K^+^ stimulation, we used an antibody against myosin-II (which recognizes the heavy chain Zipper - Zip) (**Fig. 3 a**) and two protein traps for myosin-II, GFP tagged Spaghetti Squash - Sqh (regulatory light chain) and Zip (heavy chain) (**Supplementary Fig. 6b**). Myosin-II staining was identical when using either the antibody or protein traps, reinsuring that it represents the endogenous protein localization. We observed that myosin-II is localized in MN terminal, sometimes visibly outlining the boutons, but is also very abundant in muscle and trachea. (**Fig. 3 a** and **Supplementary Fig. 6c**). Interestingly, upon stimulation, myosin-II clearly becomes enriched in new boutons, with noticeable accumulations at the base or at the edge (**Fig. 3a’**). This distribution is compatible with myosin-II function in bouton initiation and/or stabilization/retraction.

**Fig.3:**
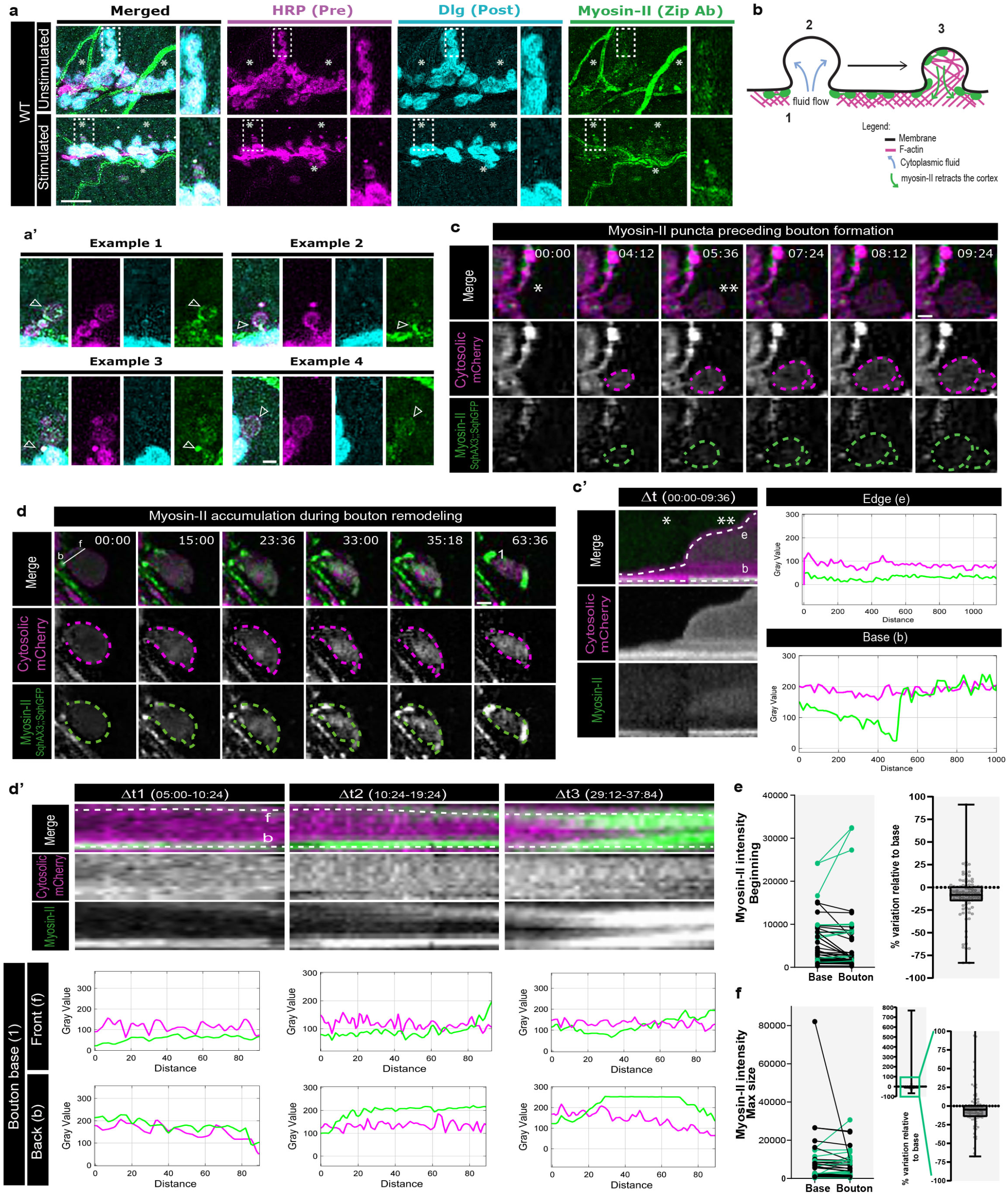
Myosin-II, a master regulator of blebbing, is dynamically recruted to new boutons. **a,** Antibody staining for myosin-ll (anti-Zipper, green) in WT (*w1118*) to see endogenous distribution at the NMJ. Unstimulated NMJ (top). Stimulated NMJ (bottom). Presynaptic membrane labeled with HRP (magenta) and postsynaptic membrane labeled with Dig (cyan). New boutons (asterisk) were identified by the lack of Dig. Scale bar, 10 μM. Myosin-ll is present in the presynaptic terminal, sometimes outlining boutons, being very abundant in muscle and trachea, n? 6 larvae, >28 NMJs and 3 biologically independent experiments, a’, post-stimulation myosin-ll is recruited to new boutons with visible accumulations at bouton base and edge (white arrows), **b**, Schematics of myosin-ll localization during bleb phases, c,d, Time-lapse images of myosin-ll dynamics during bouton formation. Bouton gorwth was visualized with cytosolic LexAop-mCherry under the control of LexA driver DvGlut and to label myosin-ll we used GFP-tagged Sqh (light chain) in a Sqh-null background (SqhAX3). Scale bar, 2 μM. Asterisk indicates where bouton will emerge, **c**, Example of myosin-ll puncta at the base preceding bouton formation, **d**, Example of myosin-ll accumulation during bouton remodeling, **c’,d’**, Kymographs from **c** and **d**. Kymographs for distinct Δt to highlight myosin-ll dynamics. Dotted lines along the bouton edge (e) and base (b) and intensity values for membrane and myosin-ll bellow, **c**’ Myosin-ll persisted at the base during bouton growth, d’ Myosin-ll clustered at the base (represented as 1 on **d**) while bouton size was reduced (Δt2 and Δt3). We represent dotted lines along the base back and front (represented as b and f on d) and the correspondent intensity values for mCherry and myosin-ll are plotted bellow, **e,f**, At beginning myosin-ll content in boutons was lower than base, but considerably increased during bouton growth. Plots with absolute values for myosin-ll intensity (left) measured in boutons and correspondent bases at beginning (**e**) or max size (**f**). Boxplot (min to max, right) showing % of variation in myosin-ll intensity in each bouton compared to its base (used as 100% reference). Line represents the median. All data points are represented, n=9 larvae, 92 boutons.

To characterize myosin-II dynamics during activity-dependent plasticity at the NMJ, we followed both bouton growth and myosin-II dynamics in real-time. We used larvae expressing cytosolic mCherry under the control of neuronal LexA driver DVGlut and a GFP tagged Sqh in a Sqh-null background (*sqh^AX3^*) to guarantee that all myosin-II was tagged with GFP. From these movies, we observed a myriad of myosin-II dynamic behaviors that we summarize here. We observed myosin-II puncta at the base of boutons preceding formation (**Figs. 3 c,c’; Supplementary movie 6**) and myosin-II flow into new boutons after they start growing (**Supplementary Figs. 7a,a’; Supplementary movie 7**). We also show two examples of myosin-II accumulation during bouton morphological remodeling (**Figs. 3d,d’ and Supplementary Figs. 7 b,b’; Supplementary movie 8**), leading to shrinkage. Likewise, in a study using correlative video microscopy and platinum replica electron microscopy (PREM), myosin-II was reported to occur in association with actin filaments in blebs of different morphologies. Although myosin-II was enriched in late-stage blebs (with a crumple shape and dense cytoskeleton), it was also found in early-stage blebs during initial assembly of the bleb cytoskeleton, possibly to mediate contraction with the cortical layer^57^. Analogous to actin, we quantified myosin-II content in the new boutons, in the beginning and at maximum size (**Supplementary Fig. 7g**). We observed that myosin-II content in new boutons was somewhat lower than the base (70% boutons at beginning and 72% boutons at maximum size), however overall the difference was very small (median at beginning was −8% and −5% at beginning and maximum size respectively) (**Figs. 3e,f**). On the other hand, in the proportion of boutons that formed with myosin-II higher than base (~30% boutons), while at beginning we saw only small gains of myosin-II, up to 25%, when boutons reached maximum size we could detect gains up to 7.7-fold superior to 100% indicating that myosin-II was massively recruited to these boutons as they were growing. Still, our data indicates that in most of the events myosin-II was prominent at the base, which was consistent with our analysis of bouton formation dynamics and its relationship with myosin-II content (**Supplementary Figs. 7 and 8**). Moreover, myosin-II puncta, which were possibly related to myosin-II flow inside new boutons (**Supplementary Figs. 8k,l**) represented a small fraction of events (~35%). Interestingly, we found that bouton formation time was significantly faster in boutons that formed with low myosin-II (**Supplementary Fig. 8c**). This finding suggests that myosin-II recruitment to new boutons is not propelling their growth but can disturb its dynamics, by slowing growth. This is in accordance with previous studies suggesting that myosin-II recruitment, to bleb cortex drives its stabilization/retraction^64^. Altogether, our data hint that myosin-II activity can contribute to regulate activity-induced remodeling at the NMJ.

### Interfering with myosin-II alters MNs capacity to form new boutons

Myosin-II homozygous mutant embryos usually display several defects in dorsal closure, head involution and axon patterning, concomitant with myosin-II having a key role in regulating cell shape during *Drosophila* development^65^. Moreover, most severe alleles are embryonically lethal, even when complemented with other myosin mutations. Therefore, we opted to use RNAi to decrease rather than abolish myosin-II levels and assay in a cell-type specific manner myosin-II putative role for acute structural plasticity. We expressed RNAi against both the regulatory light chain, Sqh, and the heavy chain, Zip, of myosin-II under the control of the neuronal Gal4 driver NSyb to promote downregulation of this myosin motor in MNs. Surprisingly, we found that neuronal disruption of the light or the heavy chain of myosin-II increased MNs capacity to form new boutons after High-K^+^ depolarization, rather than blocking it (**Figs. 4 a,b**). Additionally, expression in MNs of a dominant negative (DN) form of Sqh – a non phosphorylatable Sqh - under the control of NSyb-Gal4, also increased activity-dependent bouton frequency (**Supplementary Figs. 9 c,d**). Overall our data indicates that reduction or inactivation of myosin-II shifted the distribution of new bouton number to increased values after High-K^+^ stimulation (**Fig. 4c** and **Supplementary Fig. 9e**), which was not seen in unstimulated preparations suggesting that these changes were not due to increased developmental bouton formation (**Fig. 4 d,e** and **Supplementary Fig. 9f,g**). On the other hand, when we expressed a constitutively active (CA) form of Sqh – a phosphomimetic Sqh – we saw no significant different in bouton production compared to control (**Supplementary Figs. 8 c-g**), which showed that a general increase in presynaptic myosin-II is not sufficient to increase bouton formation. Consistently, local effects of myosin-II inhibiting drugs have previously shown that the acto-myosin cortex acts locally to extrude blebs^61^.

**Fig.4:**
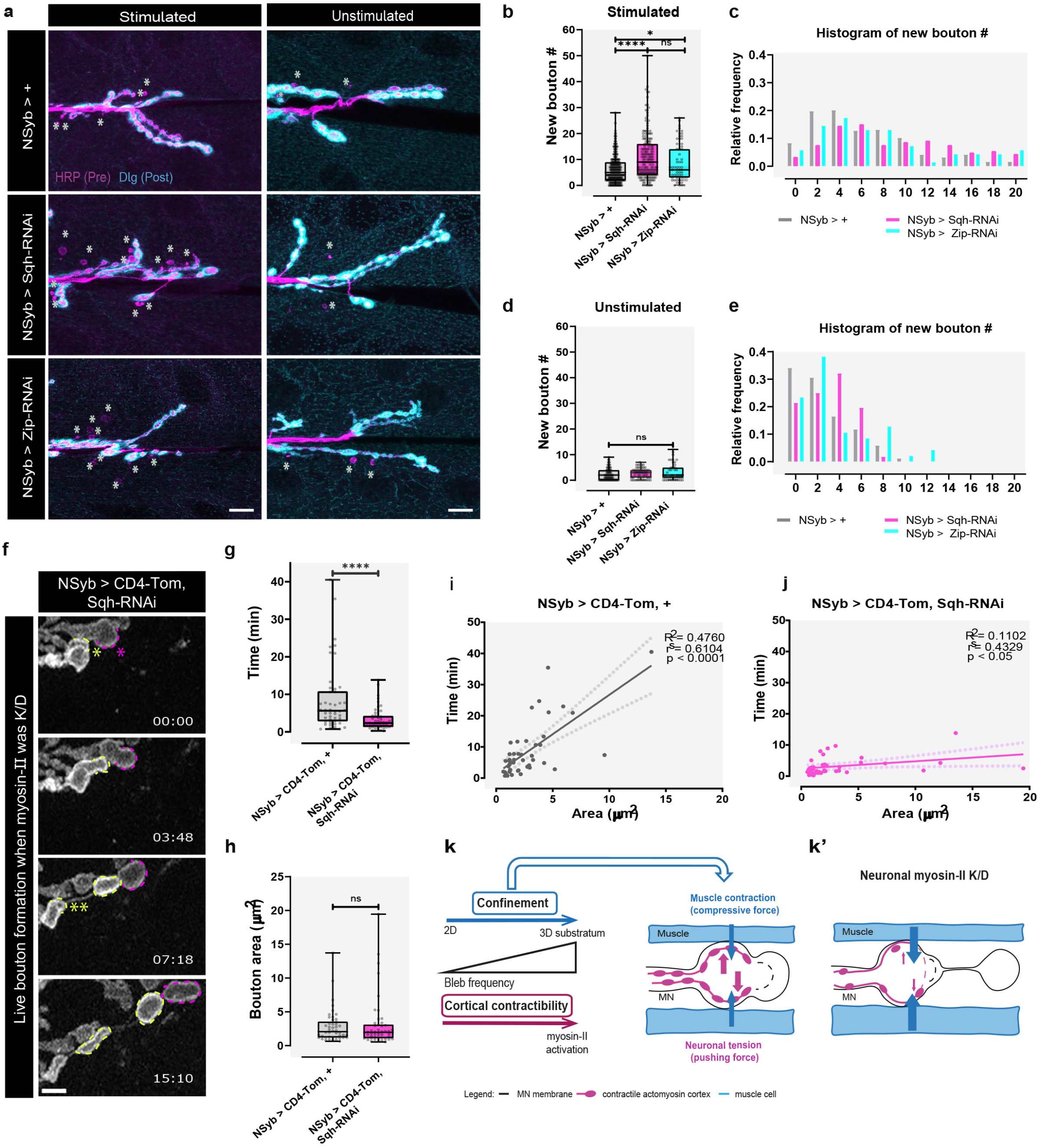
Interfering with myosin-II alters MNs capacity to form new boutons. **a,** Representative images of control and myosin-II K/D animals with (left) or without (right) High-K+ stimulation. We used NSyb-Gal4 (pan-neuronal driver) to express RNAi against two subunits of myosin-II (Sqh and Zip) in neurons. Presynaptic membrane was labeled with HRP (magenta) and postsynaptic membrane was labeled with Dig (cyan). New boutons (asterisk) were identified by the lack of Dig. Scale bar, 10 μm. **b**, Boxplot (min to max) showing that neuronal RNAi against Sqh and Zip increased new bouton frequency after acute stimulation. Line is at median. All data points are represented, **c**, Histogram showing relative frequency of new boutons in stimulated NMJs. b,c n ≥20 larvae, >69 NMJs for each line and at least 4 biologically independent experiments, **d**, Boxplot (min to max) showing that neuronal RNAi against Sqh and Zip did not change new bouton frequency during development (unstimulated control). Line is at median. All data points are represented. Statistical significance was determined with non-parametric Kruskal-Wallis test (two-tailed); ns is non-significant. **e**, Histogram showing that frequency of new boutons in unstimulated NMJs. **d, e** n ≥ 12 larvae, ≥ 47 NMJs for each line and at least 3 biologically independent experiments, **f**, Time-lapse images of bouton formation when myosin-II was K/D in neurons. In myosin-II K/D, we frequently observed threads of boutons forming; we only observed bouton formation associated with visible/intense muscle contraction. Membrane was labeled with UAS-CD4-Tom under the control of NSyb-Gal4, which also was used to drive the expression of UAS-Sqh-RNAi. Scale bar, 2 μm. Asterisk indicates where bouton will emerge, **g,** Boxplot (min to max) showing time of bouton formation is reduced in neuronal myosin-II K/D. Line is median. All data points are represented, **h,** Boxplot (min to max) showing new bouton size (area) is not altered in neuronal myosin-II K/D. Line is median. All data points are represented, **i, j,** Plots with linear regression between time and size (area) for driver control (i) and Sqh-RNAi (j). R^2^ and p-values are shown in the graphs, right on top. We also show spearman correlation values, rs, and p-value. **g-j**, n = 5 larvae, 46 boutons (for Sqh-RNAi) and 13 larvae, 46 boutons (for control), **k**, Schematics of the factors that promote bleb frequency and our hypothesis of interplay of biophysical forces: contractility mediated by myosin-II and confinement mediated by muscle contraction, **k’** Schematics of muscle contraction effect in neuronal myosin-II K/D. Statistical significance was determined with non-parametric two-tailed tests: Kruskal-Wallis (**b,d**) or Mann-Whitney (**g,h**); *p<0.05, ****p<0.0001, ns is non-significant. Significant changes observed in the distribution of new bouton number were seen across all experiments, rather in every single experiment.

At first, these results appeared unexpected, because myosin-II depletion or inhibition, strongly reduces bleb frequency in cells in 2D, but not necessarily in 3D (see below). To further analyze the dynamics of bouton formation when myosin-II is reduced in MNs, we performed live imaging of larvae with reduced myosin-II (**Fig. 4f**). We observed that when myosin-II was reduced in MNs, the NMJ was not only more plastic but boutons also formed faster (**Fig. 4g** and **Supplementary Fig. 8a; Supplementary movie 9**), while new bouton area was unchanged (**Fig. 4h** and **Supplementary Fig. 8b**). Additionally, in control animals (NSyb-Gal4>CD4Tom) the variation observed in bouton formation time was only partially explained by bouton area suggesting that other elements may regulate bouton expansion dynamics at the NMJ (**Fig. 4i**). Moreover, we could detect a clear dynamic shift in myosin-II K/D animals (**Fig. 4j**). In this case, the linear relationship was almost lost, possibly because in this scenario another factor can assume a larger effect on bouton growth dynamics. Interestingly, in our analysis of myosin-II dynamics (**Fig. 3** and **Supplementary Fig. 8e**), there were boutons that formed with low myosin-II at the base and in the bouton, a scenario of low myosin-II levels somewhat equivalent to Sqh-RNAi (despite being local), and where boutons formed faster (**Supplementary Figs. 8f,h,i**), akin to myosin-II reduction using RNAi. Strikingly, and contrary to controls (Fig.1a), in myosin-II K/D NMJs we only observed events of bouton formation concomitant with evident muscle contractions, suggesting that bouton formation in myosin-II neuronal K/D, may be dependent on muscle activity.

Overall, our data suggests that if myosin-II is necessary to initiate bouton outgrowth, its activation needs to be highly localized, like reported for blebbing cells. However, our data also provides evidence for an alternative scenario, where myosin-II is not necessary to prime the mechanism and/or to generate the cortical pressure that powers bouton growth. We suggest a model where the muscle can compress MNs creating mechanical force that can increase cortical tension in MNs to promote bouton formation (**Fig. 4k**). Remarkably, wild-type cells increase bleb frequency, and even myosin-null cells recover competence to bleb, when compressed between two agar layers^66^, which suggests that additional to internal factors, extracellular forces can also propel bleb growth. Moreover, blebs formed by cells in highly confined 3-D environments do not always sturdily depend on myosin-II contractility, supporting this hypothesis^67^. In a neuron with a weakened cortex (**Fig. 4k’**), if the initiation mechanism is primed by previous stimulation, muscle contraction can facilitate formation of new boutons from these regions. On the other hand, it is possible that in this scenario, neurons with a weakened cortex are less able to resist mechanical stresses during muscle contraction and new boutons could also arise in the weaker regions, where the cortex would break more easily. Supporting this, we frequently noticed threads of boutons forming and boutons only formed with visible/intense muscle contraction.

### Muscle contraction regulation of MN confinement is necessary for acute structural plasticity at the NMJ

Our hypothesis is that the muscle plays a mechanical role in the regulation of activity-dependent plasticity, adjusting new bouton numbers by providing different degrees of physical confinement onto MNs. To test this, our strategy was to uncouple MN excitability-muscle contraction during stimulation by either using a muscle depolarization blocker or by mechanically stretching the larvae, reducing contraction capability (**Supplementary Fig. 10**).

We used 1-Naphytlacetyl Spermine Trihydrochloride (NASPM), a Glutamate receptor (GluR) antagonist, to block muscle contraction in *Drosophila* NMJ^68^. We tested a range of NASPM (100-500 μM) to choose the lowest concentration that blocked muscle contraction to levels that did not allow the tearing of muscle tissues during the stimulation protocol. We used 300μM NASPM to test if muscle contraction contributes to activity-dependent bouton formation. After stimulation in the presence of NASPM, we observed that blocking muscle contraction had a very strong effect in bouton formation, both in wild-type and in Sqh-RNAi or Sqh-DN, suggesting bouton formation is dependent on muscle activity (**Figs. 5 d-f** and **Supplementary Figs. 11 a-c**). Moreover, blocking muscle contraction in wild-type versus myosin-II KD (or DN), eliminated completely the increase in bouton formation observed upon the reduction of myosin-II activity, supporting our hypothesis that the muscle was providing the mechanical force for bouton formation. Yet, while NASPM treatment resulted in minimal muscle contraction, it also affected postsynaptic signaling through GluRs and, consequently, muscle-MN communication, which is known to have a role in the regulation of activity-dependent bouton formation. To directly assess the effects of muscle contraction, without blocking GluRs, we mechanically stretched the larvae using extra pins to minimize muscle tearing during stimulation (**Supplementary Fig. 10).** In this situation, activity-dependent bouton frequency was significantly decreased both in Sqh-RNAi and in control NMJs (**Figs. 5g-i**); the same was observed for Sqh-DN, but not for Sqh-CA (**Supplementary Figs. d-f**).

**Fig.5:**
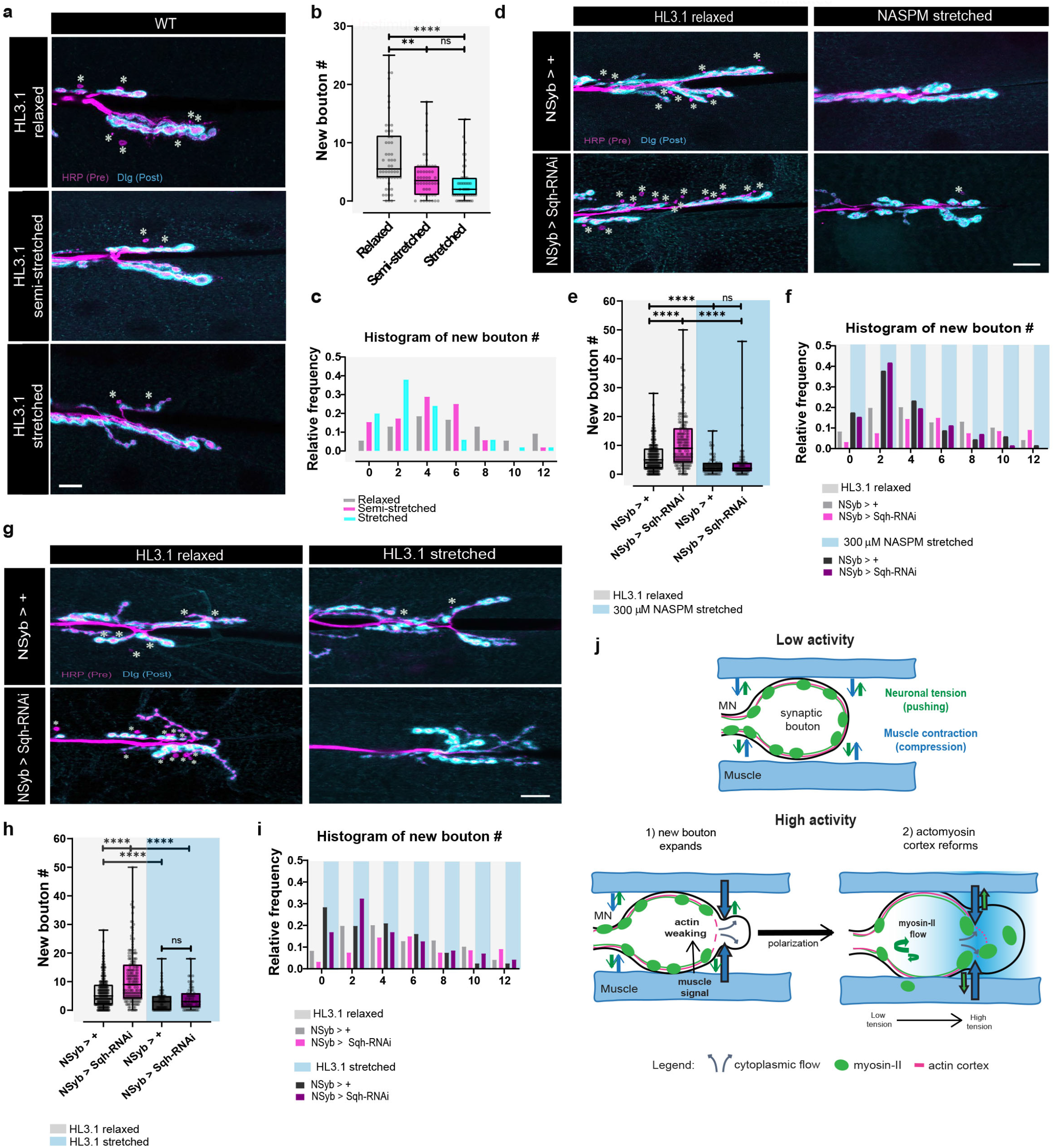
Muscle contraction regulation of MN confinement is necessary for acute structural plasticity. **a,** Images of WT (*w1118*) animals allowing distinct degrees of muscle contractibility - relaxed, semi-stretched and stretched - after High-K+ stimulation. Presynaptic membrane was labeled with HRP (magenta) and postsynaptic membrane was labeled with Dig (cyan). New boutons (asterisk) were identified by the lack of Dig. Scale bar, 10 μM. **b**, Boxplot (min to max) showing that new bouton number progressively decreased by gradually restricting muscle contractibility. Line is at median. All data points are represented, **c**, Histogram showing relative frequency of new boutons with distinct levels of muscle contractibility. **b,c** n≥8 larvae, >50 NMJs for each condition and 3 biologically independent experiments, **d**, Images of control and myosin-ll K/D animals treated with HL3.1 or 300 μM 1-Naphtylacetil spermine trihydrochloride (NASPM, GluR antagonist) after High-K+ stimulation. Presynaptic membrane was labeled with HRP (magenta) and postsynaptic membrane was labeled with Dig (cyan). New boutons (asterisk) were identified by the lack of Dig. Scale bar,10 μM. **e**, Boxplot (min to max) showing that blocking muscle depolarization and contraction with 300 μM NASPM reduced bouton formation in Sqh-RNAi and in control. Line is at median. All data points are represented, **f**, Histogram showing relative frequency of new boutons with HL3.1 or 300 μM NASPM. **e,f**, n >15 larvae, >69 NMJs for each line and 5 biologically independent experiments, **g**, Images of control and myosin-ll K/D animals where muscle contraction was either allowed, relaxed fillets, or restricted, stretched fillets, respectively, after High-K+ stimulation. Presynaptic membrane was labeled with HRP and postsynaptic membrane was labeled with Dig. New boutons (asterisk) were identified by the lack of Dig. Scale bar, 10 μm. **h**, Boxplot (min to max) showing that mechanically blocking muscle contraction by stretching the larvae reduced new bouton number in the Sqh-RNAi and in control. Line is at median. All data points are represented, **i**, Histogram showing relative frequency of new boutons for relaxed or stretched animals, n >15 larvae, >71 NMJs for each line and condition and 5 biologically independent experiments, **j**, Model for activity-dependent bouton formation at the Drosophila NMJ. MNs are deeply embebbed into the muscle, which is contractile (top). We propose that with increased activity (bottom), and in response to a not yet identified signal to induce local actin weakening, MNs add new boutons by membrane blebbing. In regions of high muscle activity, a muscle derived signal, could act to polarize boutons at these sites, and promote recruitment of molecules required/or that contribute to initiate bouton formation to spots of high membrane tension. During this process, muscle contraction can act by compressing MNs to increase their confinement, and cortical tension, and facilitate bouton formation. Statistical significance was determined with non-parametric Kruskal-Wallis test (two-tailled); ****p<0.0001, ** p<0.01, ns is non-significant.

Our results (**Figs. 4 and 5**) indicated that the increase in bouton formation observed upon myosin-II reduction or inactivation is muscle contraction-dependent and that at the *Drosophila* NMJ muscle contraction is required for normal bouton addition in response to elevated activity. Additionally, these experiments suggest that presynaptic mechanisms for force generation, such as increasing myosin-II activation in MNs, can balance with muscle contraction to modulate bouton number output in response to elevated activity. To further evaluate muscle contraction contribution for activity-dependent plasticity in the wild-type we tested the effect of distinct degrees of freedom for the muscle to contract – relaxed, semi-stretched and stretched. This treatment resulted into a correspondent gradient in bouton formation with increased number of boutons being observed with higher degree of muscle contractibility (**Figs. 5a-c**), which shows that that muscle contraction levels can strongly change structural plasticity dynamics at the NMJ. Overall, our experiments showed that muscle contraction regulation of MN confinement, an important biophysical environmental factor regulating cellular blebbing, is required to promote bouton formation at the NMJ and suggest that an intricate interplay of trans-synaptic forces may regulate activity-induced bouton number. We propose that, in addition to biochemical signaling, a balance of mechanical forces is tightly coordinated between MNs and the muscle to regulate remodeling at the NMJ – by neuronal blebbing (**Fig. 5 j)**.

## Discussion

We investigated the mechanisms of activity-dependent bouton formation *ex vivo* using the *Drosophila* NMJ, a system where intercellular and 3-D biophysical interactions are preserved. By performing high-temporal resolution live imaging, we discovered that activity-dependent bouton formation dynamics strongly resembled membrane blebbing (**Fig.1**), a mechanism widely used in 3-D migration but never reported to be used by neurons to remodel. We showed that new boutons are *bona fide* blebs, whose growth does not rely on actin polymerization but is rather pressure driven (**Fig.1–2**). Accordingly, we observed that acute bouton formation was usually correlated with muscle activity and that blocking muscle contraction significantly decreased new bouton frequency after stimulation (**Fig.5**). Our manipulation of muscle contraction resulted in predictable changes in bouton formation in response to stimulation, with a progressive increase in new bouton numbers with less stretching and higher contraction. Our results strongly suggest a mechanical role for muscle contraction in activity-dependent bouton formation. At the NMJ the MN is deeply imbedded into the muscle, which by contracting can directly alter neuronal confinement. Our hypothesis is that with elevated activity, and in response to a still unclear signal to initiate bouton growth, MNs add new boutons by membrane blebbing and muscle contraction is required to increase neuronal confinement, and consequently cortical tension, which powers bouton outgrowth. This hypothesis is in agreement with blebs being favored in tissues where cells encounter increased mechanical resistance^69^. In this scenario, as it has been proposed for cells in compressive 3-D environments^70^, it is possible that a muscle derived signal, released in regions of higher activity, can polarize the recruitment of molecules required to or that facilitate bouton formation to spots of high membrane tension.

While the molecular identity of the signal that leads to bouton initiation remains to be elucidated, it is well established that blebs nucleate when a small patch of membrane is detached from the actin cortex^70^. This can happen either as a direct consequence of buildup in hydrostatic pressure (that detaches membrane to cortex binding proteins) or by formation of local gaps, resulting from rupture of the cortex in regions of high membrane energy (where accumulation of myosin-II helps to weaken the cortex)^70^.

While the first model is adequate when cells are subjected to high compression forces, it does not perform as well when compression force is lower^70^. Our data suggests that, although at the NMJ the two mechanisms may coexist, with elevated activity boutons tend to form mainly as a consequence of compressive forces exerted by the muscle. Supporting this, we found that activity-induced bouton formation was normally rapid and correlated with visible muscle contraction, while events with low or no visible muscle contraction were slow and occasional. Importantly, blocking muscle contraction impaired activity-dependent bouton formation. Additionally, even though we observed myosin-II puncta preceding bouton formation in ~35% of events, we found boutons forming with low myosin-II levels and decreasing or inactivation of myosin-II did not prevent bouton formation. The fact that boutons always formed without filamin, which links actin to membrane, further supported that actin polymerization is not required for bouton initiation. Altogether we favor a model in which boutons form in regions of high pressure by detachment of membrane from the actin cortex and that local myosin-II activation can facilitate the weaking of the cortex at these sites.

Another question that remains unanswered is related with the membrane source required for structural changes observed in blebs, or, in our case, for neuronal remodeling. In principle, bleb expansion can be sustained by unfolding of local membrane invaginations or by localized exocytosis at the base of the bouton^54^. It has been shown that blebs can form by unfolding of local intracellular membrane invaginations^71^, but it is not known whether this occurs in neurons, or whether local membrane addition is needed. Our live imaging with CD4-Tom revealed local intracellular vesicle dynamics in MNs at the place of bouton formation. Interestingly, it was found that intense activity *in vivo*, such as locomotion or seizures, also induces rapid formation of new boutons, a process that depends on Synapsin activation by PKA^39^. These new boutons were filled with SVs and showed several profiles of SVs docked and partially fused with the membrane, suggestive of exocytic events, but the EM tomographic analysis also showed the presence of membrane folds within the new bouton^39^. In conclusion, there is data supporting both models for membrane expansion during blebbing, and more experiments will be necessary to clarify this question. Furthermore, SVs are enriched in Synapsin, a protein that has been suggested to organize SVs into a liquid phase in which SVs remain tightly clustered, forming a dense compartment^72^, and we suggest that this increased density could potentially also help to generate local cytosolic pressure during bouton outgrowth.

Synaptic boutons have been consistently associated with axonal GCs and actin mediated protrusive processes, such as lamellipodia or filopodia, which are typically seen in cells migrating in 2-D. However, *in vivo*, cells adjust their shapes through complex 3-D microenvironments, and are subjected to varying degrees of physical confinement provided by neighboring cells and tissues^69,73^. In fact, cells migrating in confined environments typically display rounder morphologies and use hydrostatic blebs for movement. Interestingly, a recent study showed that neurons from mouse central nervous system (CNS) growing in 3-D matrixes displayed GCs that had an amoeboid-like morphology, characterized by very low adhesive interactions and extensive rounded deformations of the bod wall^74^, challenging the long-lasting paradigm of the lamellipodia-base GC. In the brain, even though there are no muscle compressive forces, synaptic boutons are equally confined surrounded by other neuronal, astrocytic or microglial processes. Interestingly, astrocytes and microglia have been shown to regulate many aspects of neuronal function, including plasticity^75–77^, with glia having dynamic protrusions that directly contact synapses. Here, we put forward the hypothesis that both astrocytes or microglia could potentially provide the physical confinement, required to facilitate synaptic bouton formation, or even help to specify the blebbing sites, but this model will need to be tested. On the other hand, it will be interesting to study how neurons can sense and respond to mechanical signals provided by the cellular environment, and if pathways used for mechanosensing and transduction, likely regulating adhesion and contacts with extracellular partners, can be coupled with synaptic transmission machinery during plasticity.

Our model explains the formation of new synaptic boutons by a physical process, which in addition to the more studied transcriptional or biochemical factors, allows for a fast, on demand expansion of the NMJ. Without a bleb-like mechanism, we would predict a slower NMJ expansion, not always compatible with the requirements of the neuromuscular system during development or increased activity. Whether synapses are always susceptible to bleb-dependent expansion or require priming by cytoskeletal regulation remains to be elucidated. Moreover, considering the conservation of presynaptic cytoskeletal components and bouton ultrastructure throughout evolution^6^, we postulate that this mechanism of synapse remodeling can be present in other organisms, including vertebrates. Overall, our finding that bouton addition at the NMJ occurs as result of a MN-muscle physical interplay highlights the importance of intercellular cooperation during plastic changes. Circuit remodeling as a response to experience requires rapid modulations of bouton number^78^. We speculate that the regulation of confinement by non-neuronal cells (muscle or glia) can be a mechanism widely used by the nervous system to coordinate local activity-dependent structural changes in neurons with its surrounding 3-D cellular microenvironment. Future studies will elucidate whether the understanding of the biochemical and mechanical relations between these cells during structural plasticity can help design strategies to ameliorate MN loss or dysfunction present in many neuromuscular diseases. Additionally, we also establish the genetically tractable *Drosophila* NMJ as an *ex vivo* system in which to study blebbing in 3-D, with enormous potential to manipulate and visualize each of the cell types.

## Methods

### Drosophila culture and stocks

*Drosophila melanogaster* stocks were maintained at 25°C on standard media. Crosses were set with the minimum of 5-10 virgin females and 3-5 males of the appropriate genotype. Adults were removed 7-8 days after each cross to ensure segregated generations. For larval collection, eggs were laid and grown on apple juice vials at 25°C. For RNAi experiments, collection of RNAis expressing strains and their controls were set up at 29-30°C to maximize the efficiency of knockdown. TM6b or GFP expressing balancer chromosomes were used to facilitate genotyping of larva (non-tubby or non-fluorescent larvae were selected respectively). The *w^1118^* line was used as the wild-type (WT) genotype, as the stocks used are in this genetic background. For tissue-specific transgene expression NSyb-Gal4 (Pan-neuronal driver) was used to drive UAS-constructs’ expression in neurons. UAS lines, mutant alleles and recombinant used are describe in the tables below. As driver control, we used nSyb-Gal4 crossed with *w^1118^*.

**Table 1.** UAS and Gal4 lines used:

| Name | Description | Reference |
| --- | --- | --- |
| UAS-Zip-RNAi | Expresses RNAi against Zipper (heavy chain of NMMII) under the control of UAS; chromosome 3 | BDSC #36727 |
| UAS-Sqh-RNAi | Expresses RNAi against Spaghetti squash (heavy chain of NMMII) under the control of UAS; chromosome 3 | BDSC #32439 |
| UAS-Sqh-CA | Expresses a phosphomimic Sqh protein with T20E and S21E aa substitutions under the control of UAS; chromosome 3 | BDSC #64411 |
| UAS-Sqh-DN | Expresses a non-phosphorylatable Sqh protein with T20A and S21A aa substitutions under the control of UAS; chromosome 3 | BDSC #64114 |
| NSyb-Gal4 | Pan-neuronal driver neuronal synaptobrevin-Gal4 | Tom Schwarz Lab |

\*BDSC -Bloomington Stock Center

**Table 2.**
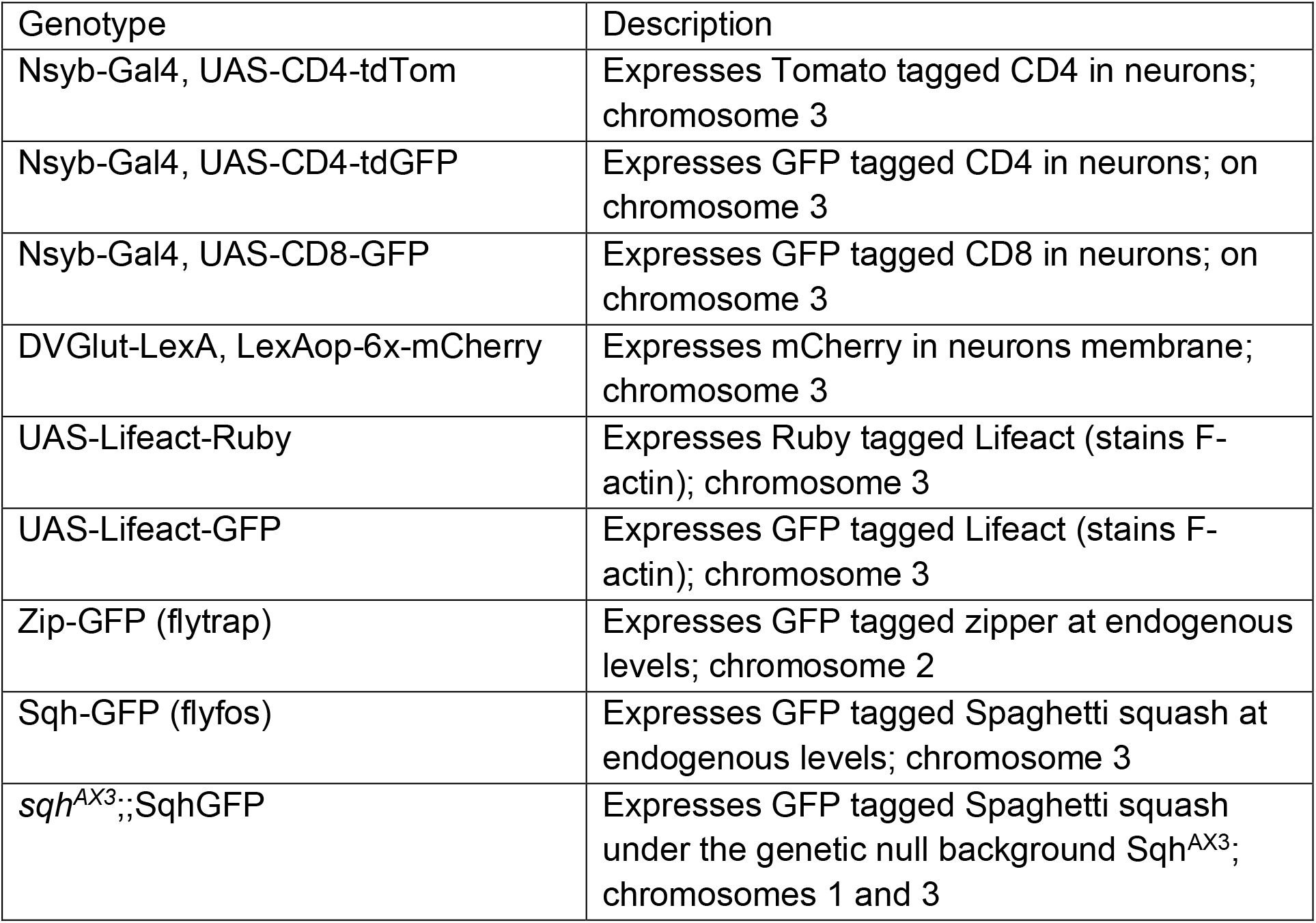
Recombinants and other lines:

### Generation of transgenic flies

The DVGlut-LexA^79^ driver was derived from the pCaSpeR VGlut >> LexA construct^80^ by removing the FRT cassette with KpnI restriction enzymes. Transgenic lines were generated by standard P-element-mediated transformation procedures in a yw background. Random insertions were selected based on strength and specificity. 13XLexAop-6XmCherry-HA (attP2) from Bloomington, BL52271.

### Larval dissections and immunocytochemistry

Third instar larvae were dissected in HL3.1 saline (in mM: 70 NaCl, 5 KCl, 0.1 CaCl2, 4 MgCl2, 10 NaHCO3, 5 Trehalose, 115 Sucrose, 5 HEPES-NaOH, pH 7.3-7.4) using a procedure similar to Brent *et. (*2009). Gut and fat body were removed, while the CNS was kept intact until after fixation. The resulting larval fillets were fixed in Bouin’s fixative (saturated picric acid + formaldehyde + glacial acetic acid) or in PFA (4% paraformaldehyde diluted in 1x PBS) at room temperature for 5 and 20min, respectively, then extensively washed in PBT (1x PBS + 0,1-0,3%Triton) to permeabilize membranes. Blocking of unspecific binding was done incubating 30 min-1 hour with NGS (Normal Goat Serum) or BSA (Bovine Serum Albumin) dissolved in PBT. Primary antibody incubation was performed overnight at 4°C, in blocking solution. Subsequently, larvae were extensively washed using PBT, followed by blocking for 30min-1h and incubated for 2 hours with secondary antibody at room temperature. After extensive washing using PBT larvae were transferred to 50% glycerol in PBS for 5 min and then mounted in DABCO medium (in 100% glycerol).

**Table 3.** Primary antibodies used in this study:

| Antigen and clone ID | Species | Concentration | Fixative | Reference/Source |
| --- | --- | --- | --- | --- |
| Dlg(4F3) | Mouse | 1:250 | Bouin's or PFA | Hybridoma bank |
| Dlg | Rabbit | 1:10000 | PFA | Gift from Vivian Budnik |
| GFP | Rabbit | 1:1000 | PFA | Life technologies A11122 |
| NM-II (heavy chain) | Rabbit | 1:1000 | PFA | Gift from Christine Field |
| Filamin (N terminus/aa 189-482) | Rat | 1:800 | Bouin's | (Külshammer and Uhlirova, 2013)/ Gift from M. Uhlirova Lab |

All secondary antibodies used in this study were purchased from Jackson Immunoresearch and used at 1:500 (initial dilution in 50% glycerol, according to manufacturer indications). Also from Jackson Immunoresearch Horseradish peroxidase (HRP) conjugated to Cy3, Alexa488 or Alexa647 was used to label neuronal membrane (1:500).

### High-K^+^ stimulation of larval NMJs

Third instar larvae were pinned onto Sylgard-coated plates using insect pins and partially dissected in HL3.1 saline solution (in mM: 70 NaCl, 5 KCl, 0.1 CaCl2, 4 MgCl2, 10 NaHCO3, 5 Trehalose, 115 Sucrose, 5 HEPES-NaOH, pH 7.3-7.4) at room temperature. Importantly, prior stimulation the dissection pins were moved inward to the same guide shape at ~50% of the original size of each larva to allow for muscle contraction. We used the stimulation protocol described by Vasin *et*. (2015) that uses a spaced High-K^+^ depolarization paradigm to induce patterned activity and rapid bouton formation at *Drosophila* larval NMJ. Briefly, relaxed fillets were subjected to incubations in High K^+^ (90 mM) and High Ca^2+^ (1.5 mM) HL3.1 adjusted for osmolarity changes with 2, 2, 2 min pulses each separated by 10 min incubation in normal HL3.1. For immunocytochemistry, stimulated larvae were fixed 30 min after the 3^rd^ stimulus to maximize bouton formation in the respective paradigms. Control larvae (nonstimulated) are dissected as described above but with only normal HL3.1. The protocol for acute stimulation of boutons used for live imaging was identical the one described above, unless otherwise stated. Additionally, for live imaging, we also used a mass stimulation procedure of 10- or 16-min High-K^+^ incubations developed by Martin *et*. (2017). Briefly, third instar larvae are dissected and glued directly onto a Sylgard-coated slide and imaged in HL3.1 saline right after stimulation for 30 min to 1.5 hours. The ventral nerve cord and the CNS were maintained intact and stretching was minimized as much as possible.

### Drug administration and manipulation of muscle contractibility

#### Actin manipulation

actin depolymerizer Latrunculin B (Focus Biomolecules) and actin stabilizer Jasplakinolide (ChemCruz) were used. These reagents were prepared as 25 mM and 3 mM stocks in DMSO respectively and diluted in HL3.1 and in High-K^+^ High-Ca^2+^ HL3.1 solutions to the desired concentration. Drug treatments were performed by pretreating dissected larval preparations in HL3.1 solution containing 10 μM Latrunculin A or 10 μM Jasplakinolide for 15 min. Stimulation was performed using HL3.1 and High-K^+^ High-Ca^2+^ HL3.1 solutions containing either 10 μM Latrunculin B or 10 μM Jasplakinolide. Each experiment was always performed side-by-side with a normal stimulation control, using HL3.1, and a solvent control, using DMSO (dilution used for LatB or JAS from stock).

#### Muscle inactivation

to prevent muscle contraction, we prepared a 100 mM solution of 1-Naphtylacetil spermine trihydrochloride (NASPM) in H20, which was diluted to the desired concentration (mostly 300 μM NASPM solution) in HL3.1 and the same was done for High-K^+^. The solution HL3.1 containing NASPM was incubated with the preparation in the dark for 30 min prior to stimulation protocol. We also used NASPM (Sigma, N193) during the stimulation protocol. During stimulation the microscope light was turned off to minimize light exposure to the drug and we maintained the larva fully stretched with 6 pins to minimize residual contraction to verify drug efficacy (NMJs that contracted significantly exhibited visible muscle damage or tearing). To directly assess the role of muscle contraction during bouton formation larvae were mechanically stretched and incubated with HL3.1 only (30’ pre-treatment and during stimulation). To block muscle contraction and simultaneously avoid preparation tearing larvae we stretched with 10 pins (more 4 pins were placed surrounding the NMJ 6/7 between A2-A4). To minimize protocol variations related with room temperature and/or stimulation solution pH, each experiment (for NASPM and mechanical stretching) was always performed side-by-side with a normal stimulation control, with stimulation starting after a mock 30’ treatment in HL3.1 only. Additionally, to test if diverse degrees of contractility can result in discrete plasticity outputs larvae were divided into 3 groups: relaxed (2 pins, half size), semi-stretched (2 pins, full size) and stretched (6 pins not fully stretched to avoid muscle tearing). In this case, the relaxed larvae were used as normal stimulation control.

### Fluorescence imaging

Confocal images were obtained on a Laser scanning confocal microscope (LSM 710) with a 40x 1.3 NA water-immersion objective or a 63x 1.3NA oil-emersion objectives (Carl Zeiss). Images were processed in image J (National Institutes of Health) and Adobe Illustrator or Photoshop software. The live-imaging experiments were performed with a spinning Disk confocal microscope (Andor) using a 60X 1.3 NA oil immersion objective (Carl Zeiss); equipped with a heating stage heated to 25°C. Quantification of bouton number was performed at NMJ 6/7, abdominal segments A2-A4 were analyzed. In general, at least 12 (fixed) or 10 (live) NMJs of each genotype were analyzed for each experiment/time-point. Quantitative and video analysis was performed using maximum intensity projections from z-stacks on image. Images were mounted in Adobe Illustrator and Photoshop.

### Kymographs analysis

Kymographs were used to visualize motion of actin or myosin-II between the base and edge of new boutons and extract their dynamic behaviour during live bouton formation. Since our movies show a lot of movement (due to contractions of larval preparation) prior to generation of kymographs, videos were divided into discrete smaller Δt’s. In the cases in which this strategy was not sufficient to remove the movement, we also corrected by rigid registration (2d/3d + t) in ImageJ/Jiji; membrane channel was aligned or adjusted to actin or myosin-II. Kymographs were build using Multi Kymograph macro from ImageJ/Fiji. We draw a line ROI from the base that passed through the bouton reaching the edge and plotting this profile, encoding the intensity in grayscale level (or color for color images) for all slices in the Δt. To adjust the line position, we select a frame where the bouton is still small and draw a line ROI preferentially from the center through the bouton. However, sometimes this was not possible. In these cases, we used the max size frame as reference and draw a line ROI that extended from base to edge during bouton growth. In the kymographs, as they are oriented in the figures, the x-axis reports time (from beginning) and y-axis reports distance (from base). After kymographs were generated, to measure changes that occurred at the base or at the edge, we traced for membrane and actin/myosin-II a line along the base or the edge and plotted the fluorescence intensity values for these lines. In these plots, x-axis reports time (along the line that we draw) and y-axis is fluorescence intensity along this time window.

### Analysis of fluorescence intensity

For analyzing fluorescence intensity of actin or myosin-II, maximum intensity projections from z-stacks were used. For each event the intensity (integrated density) was measured in a 1×1 μm square ROI at the base (beneath the growing bouton) and in the bouton (near the edge); in both cases intensity was measure at the initial frame (when bouton starts to emerge) and at maximal size (when bouton stopped growing). To discount for photobleaching effects we also measured intensity of fluorescence at main branch of the NMJ in the initial frame and when bouton reached max size. We used a 1×1 μm square as ROI. To quantify actin or myosin-II content we subtracted intensity measured in bouton to the intensity measured in the base (base – bouton). If base-bouton= 0, bouton equal to base; if base-bouton > 0, bouton lower than base; if base-bouton < 0, bouton higher than base. The % of variation inside boutons was calculated using the actin or myosin-II at the base as 100% reference; the value was corrected for photobleaching effects by subtracting % of change observed in the main branch (intensity at initial frame – intensity at maxsize). To quantify actin or myosin-II fluxes were measured by subtracting intensity measured in max size at to the intensity measured in the beginning (beginning – max size). If beginning-max size=0, max size = beginning; if beginning – max size > 0, max size < beginning; if beginning – max size < 0, max size > beginning. The % of variation was calculated using the actin or myosin-II at maxsize as 100% reference; the value was corrected for photobleaching effect by subtracting % of change observed in the main branch (intensity at initial frame – intensity at intensity at maxsize).

### Statistical analysis

Statistical analyses were conducted using GraphPad Prism Version 6.0 (GraphPad Software).

First, we performed a descriptive analysis of the data to study the distribution and variance associated with each condition/genotype sampled. Furthermore, all data sets were tested for normality using Shapiro-Wilks normality test. We opted to perform a non-parametric based inferential statistical analysis since we found the data to be asymmetrically distributed, high covariance levels (superior to 60%) and frequently data sets did not pass normality. Statistical significance in two-way comparisons was determined by Mann-Whitney test, while Kruskal-Wallis analysis was used when comparing more than two datasets. In both cases we performed two-tailed tests. In all figures data is presented as median (interquartile range); ****p<0.0001;***p<0.001; **p<0.01; *p<0.05, n.s. not significant. Statistical comparison is made between all groups. Sample size is presented in the Table 4.

**Table 4.**
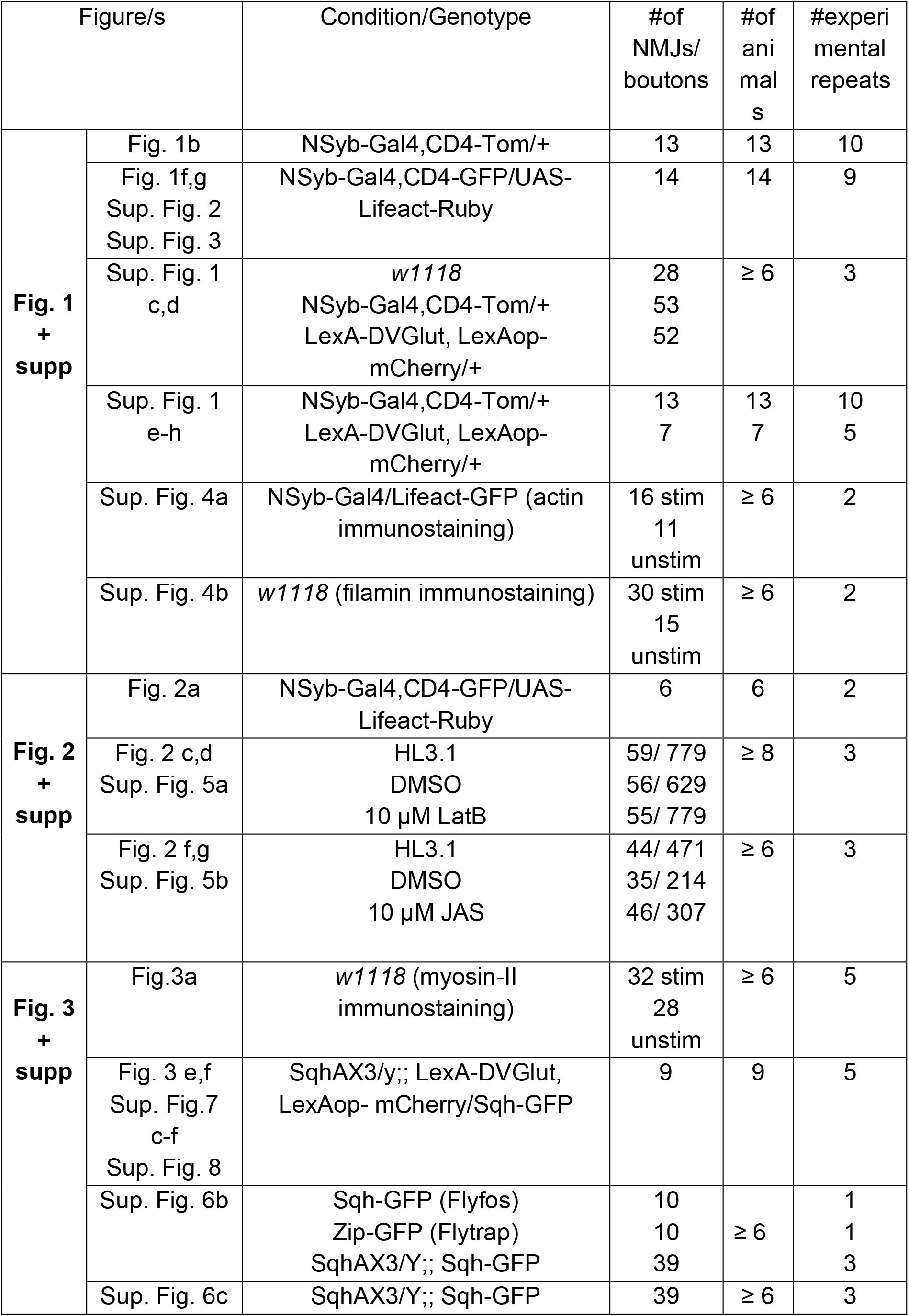
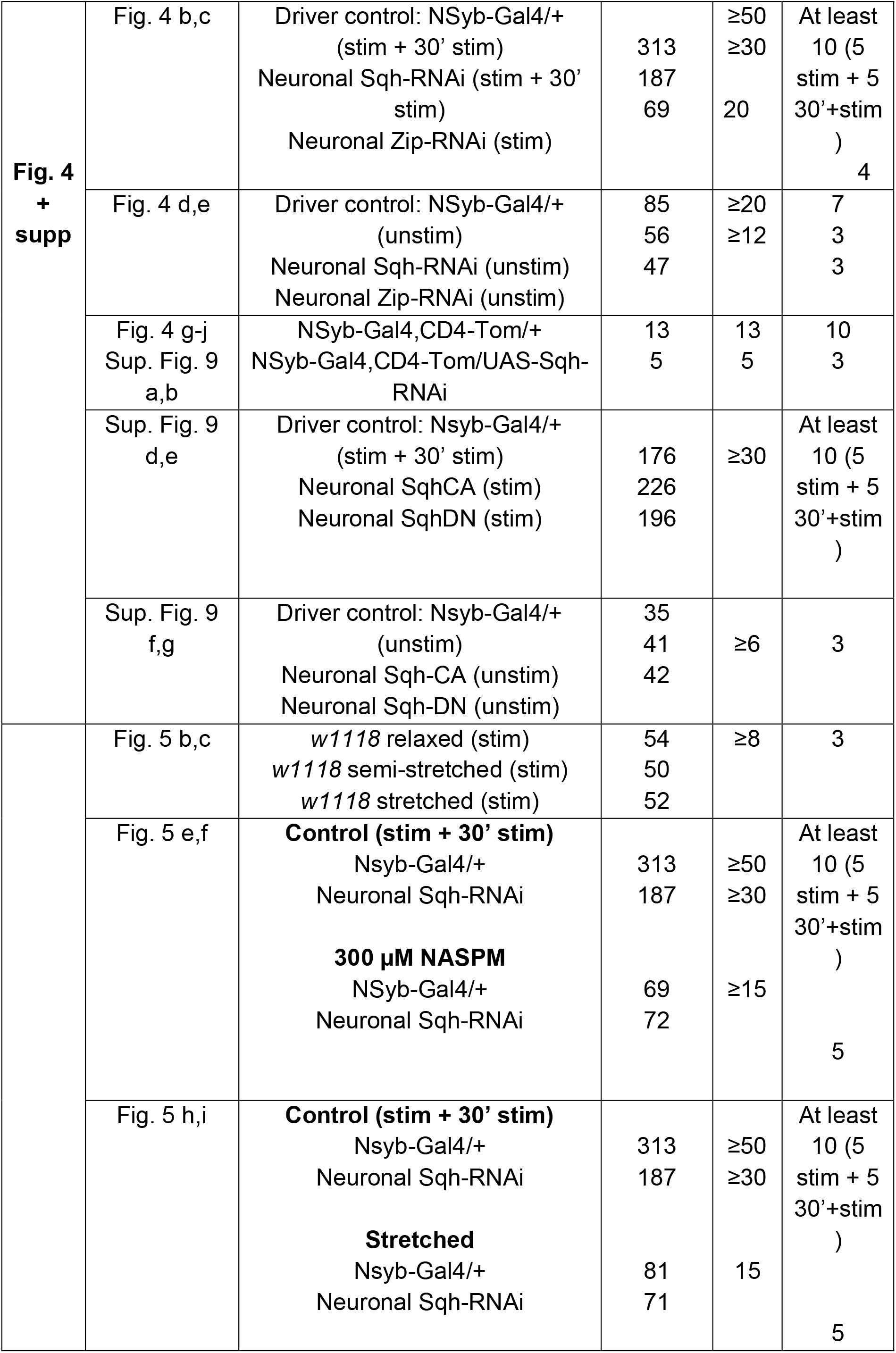
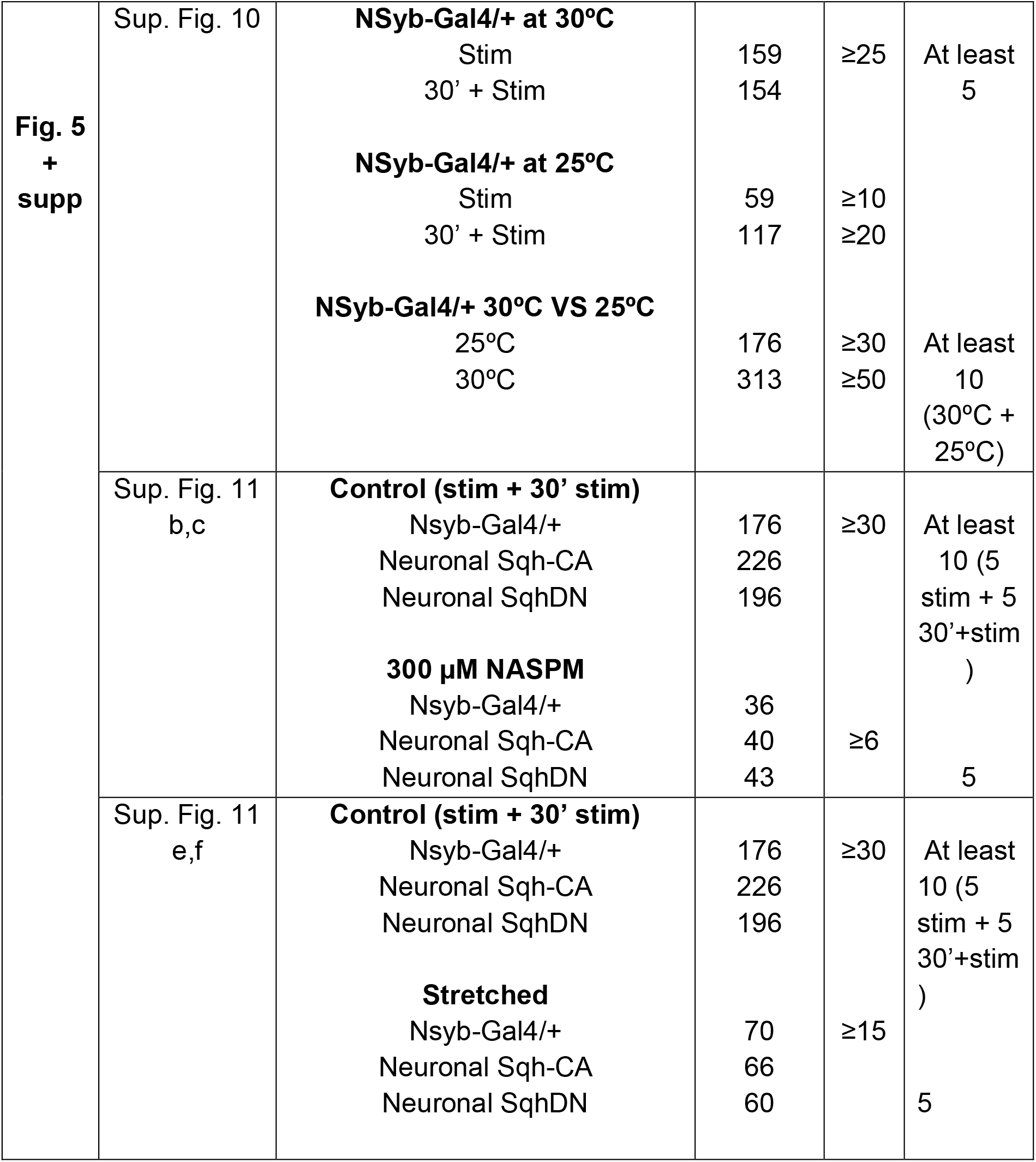
Number of animals used in quantified NMJ data.

## Data availability

Data supporting the findings of this study are available from the corresponding author on request. Source data are provided with this paper.

## Acknowledgements

We would like to thank Telmo Pereira from the Microscopy Facility for technical support, the Fly Facility at CEDOC; CONGENTO: consortium for genetically tractable organisms. We thank the Developmental Studies Hybridoma Bank, and Bloomington Drosophila Stock Center for antibodies and fly stocks. This work was supported by H2020 Marie Skłodowska-Curie Actions [H2020-GA661543-Neuronal Trafficking and 752891, GEMiNI to R.O.T. and C.S.M., respectively], Fundo Regional para a Ciência e Tecnologia [IF/00392/2013/CP1192/CT0002 and IF/01154/2014/CP1252/CT0003 to R.O.T. and C.S.M., respectively]. A.R.F. is supported with a PhD scholarship from Fundação para a Ciência e Tecnologia, Portugal, with the reference SFRH/BD/144488/2019. This work was also supported by iNOVA4Health – UIDB/04462/2020.

## Author contributions

A.R.F and R.O.T designed and executed the experiments. A.R.F. analyzed the data. A.R.F. And R.O.T. wrote the manuscript. C.S.M. built the LexA-DVGlut driver, contributed with intellectual input and edited the manuscript. E.R.G. contributed with intellectual input and edited the manuscript.

## Materials & Correspondence

Correspondence and material requests should be addressed to A.R.F or R.O.T. Emails for contact are, respectively.

## List of supplementary figures

Supp. Fig.1 - Effect of membrane tags on bouton formation frequency and dynamics.

Supp. Fig.2 - Quantification of actin content in new boutons and bouton formation dynamics.

Supp. Fig.3 - Bouton classes based on actin changes in bouton and base and bouton formation dynamics.

Supp. Fig. 4 - F-actin and filamin localization at the NMJ before and after High-K^+^ stimulation.

Supp. Fig. 5 - New bouton number after manipulation of actin dynamics.

Supp. Fig. 6 - Myosin-II-protein trap localization at the NMJ after High-K^+^ stimulation.

Supp. Fig. 7- Quantifications of myosin-II content in bouton and bouton formation dynamics.

Supp. Fig. 8 - Bouton classes based on myosin-II changes in bouton and base and bouton formation dynamics

Supp. Fig. 9 - Effects of reducing myosin-II or altering activity of myosin-II in activity-dependent plasticity.

Supp Fig. 10-Experimental design for blocking muscle contractions during activity-dependent plasticity and controls

Supp. Fig. 11-Effects of blocking muscle activation and/or contraction in activity-dependent bouton formation with Myosin-II activation or inactivation.

## List of supplementary movies

Supp. movie 1 – Bouton formation in WT NMJ: Fast bouton formation (with muscle contraction)

Supp. movie 2 – Bouton formation in WT NMJ: Slow bouton formation

Supp. movie 3 – Bouton showing bleb life phases

Supp. movie 4 – Sequential bouton formation without retraction

Supp. movie 5 – Sequential bouton formation with retraction

Supp. movie 6 – Myosin-II puncta preceding bouton formation

Supp. movie 7 – Myosin-II flow during bouton expansion

Supp. movie 8 – Myosin-II accumulation during bouton remodeling

Supp. movie 9 – Bouton formation in neuronal myosin-II K/D NMJ

