## Supplementary material for "Motor neuron boutons remodel through membrane blebbing": Suplemmentary Figures 1-11

### Supplementary Fig. 1

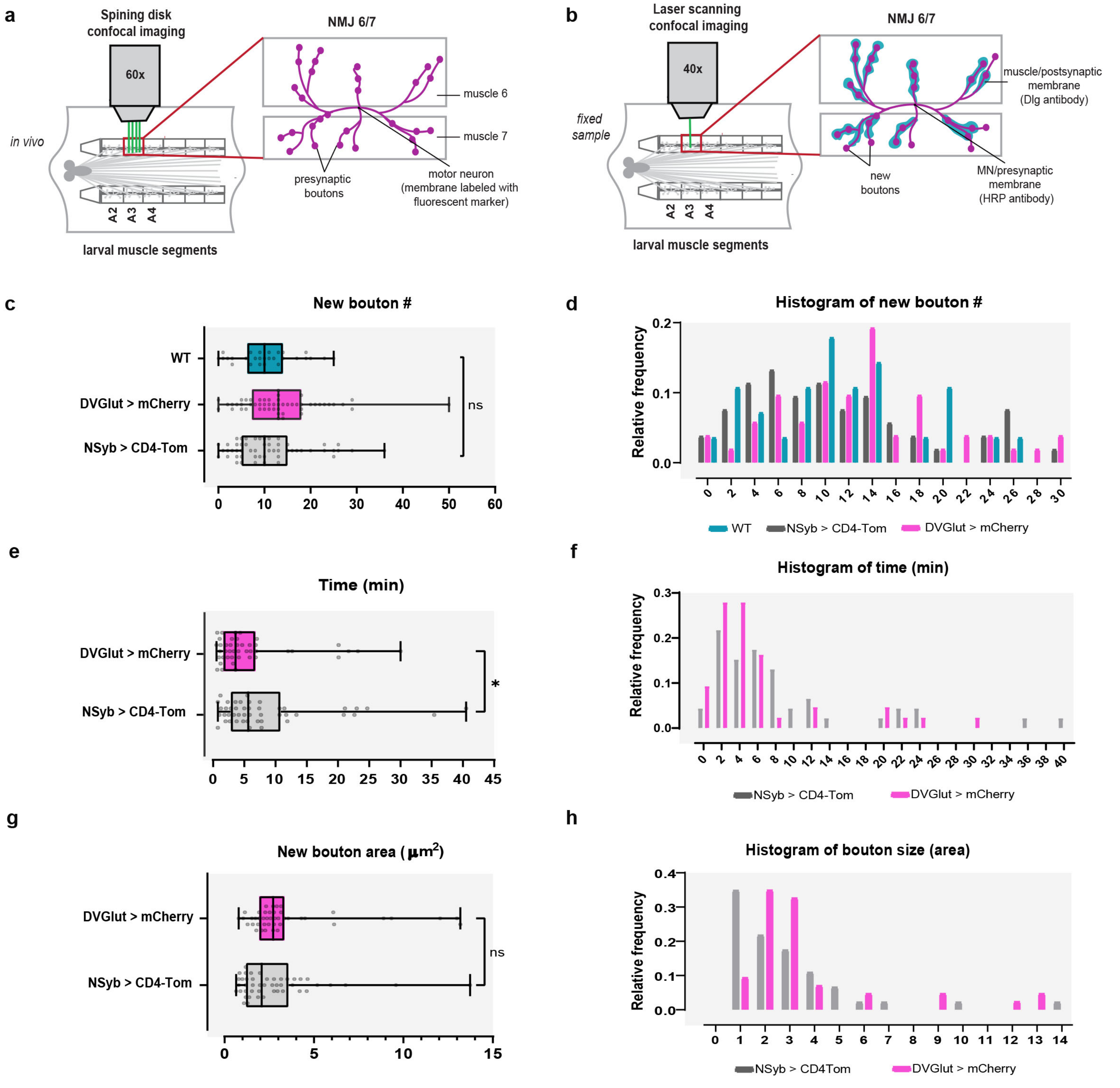


**Supplementary Fig. 1 Effect of membrane tags on bouton formation frequency and dynamics. a, b** Schematics of experimental setup, showing the dissected larval preparation and the NMJs imaged in this manuscript. **a,** Spinning disk confocal imaging allowed to follow in vivo and in real time bouton growth. Animals expressing fluorescent tags – CD4-Tomato (CD4-Tom, transmembranar) and mCherry (mCherry, cytosolic), under the control of NSyb-Gal4 and LexA Driver DVGlut, were used to visualize neuronal membrane. **b,** Laser scanning confocal imaging was used to quantify new bouton number in fixed samples. Presynaptic membrane was labeled with HRP and postsynaptic membrane was labeled with Dlg. New boutons were identified by the lack of Dlg. **c,** Boxplot (min to max) showing that new bouton frequency was unaltered when comparing WT animals (*w1118*), with neurons expressing CD4-Tom or mCherry. Line represents the median. All data points are represented. **d,** Histogram showing relative frequency of new boutons for WT and membrane tags. n ≥ 6 larvae, n ≥ 28 NMJs for each line and 3 biologically independent experiments. **e,** Boxplot (min to max) showing time of bouton formation is increased by using CD4-Tom compared with mCherry. Line is median. All data points are represented. **f,** Histogram showing relative frequency of times of bouton formation for membrane tags. **g,** Boxplot (min to max) showing new bouton size is unchanged when using CD4-Tom or mCherry. Line is median. All data points are represented. **e-g,** n = 13 larvae, 46 boutons (for CD4-Tom) and 7 larvae, 43 boutons (for mCherry). Statistical significance was determined with two-tailed non-parametric tests: Kruskal-Wallis test (**c**) or with Mann-Whitney test (**e,g**); *p<0.05, ns is non-significant.

Suplementary Fig. 2

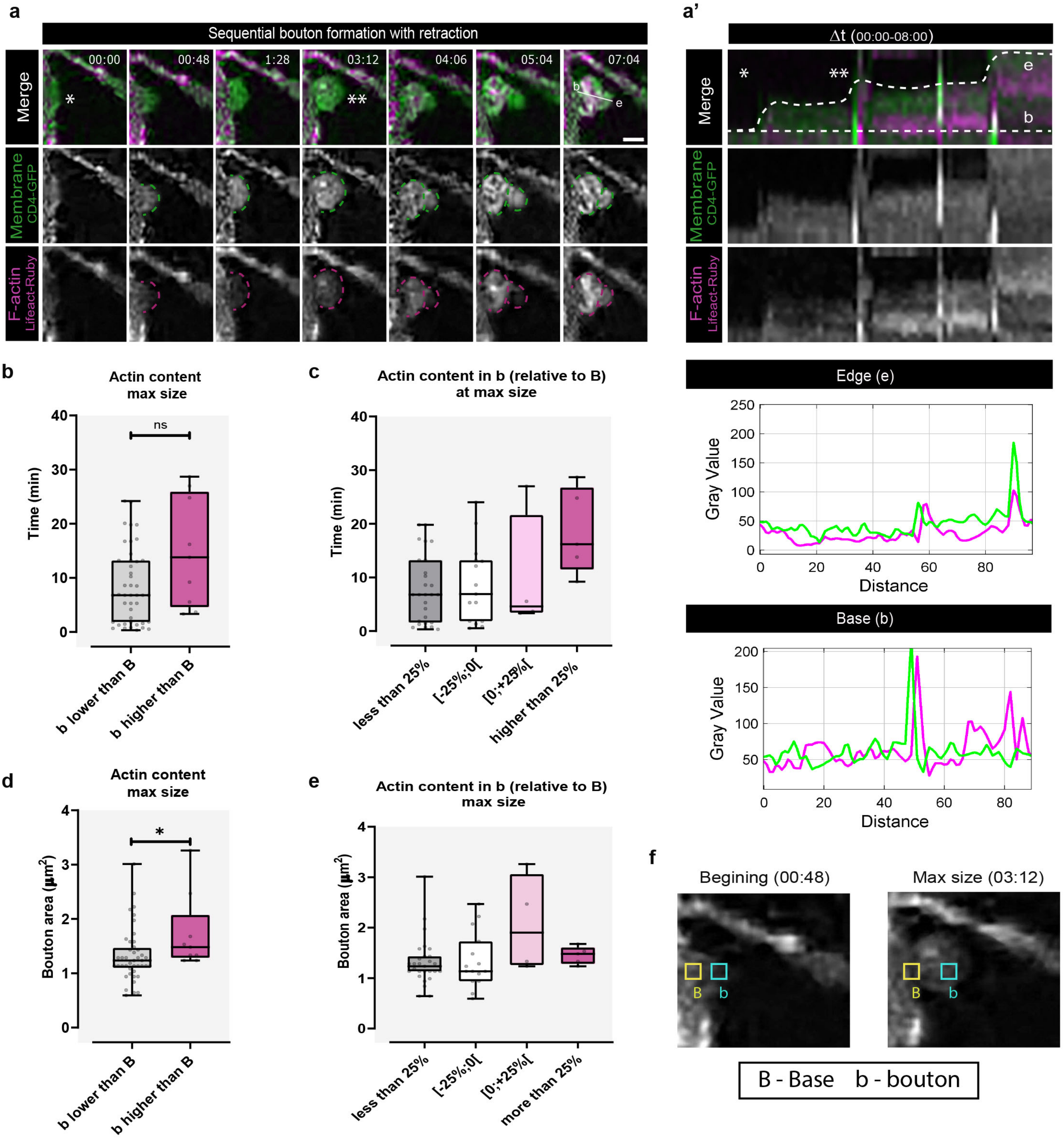


**Supplementary Fig. 2 Quantification of actin content in new boutons and bouton formation dynamics.** a, Time-lapse image of sequential bouton formation with retraction. Neuronal membrane and F-actin were labeled with UAS-CD4-GFP with UAS- Lifeact-Ruby, both under the control of NSyb-Gal4. Scale bar, 2 μm. Asterisk indicates where bouton will emerge. **a',** Kymographs from a showing that in the beginning actin is weak and accumulates when boutons stop growing/retract. We represent dotted lines along the bouton edge (e) and base (b) and the correspondent intensity values for membrane and F-actin are plotted below. **b,** Boxplot (min to max) showing that times of bouton formation were unchanged in boutons (b) with actin lower or higher than base (B). Line is median. All data points are represented. **c,** Boxplot (min to max) showing times of bouton formation in bouton classes based on actin content relative to base: less than 25%, [ -25%; 0[, [0; +25% [, more than 25%. Line represents the median. We noticed a non-statistical trend towards increased bouton formation time with higher actin content. All data points are represented. **d,** Boxplot (min to max) showing that bouton area was increased in boutons (b) in which actin was higher than base (B), compared with boutons that had actin lower than the base. Line is median. All data points are represented. **e,** Boxplot (min to max) showing bouton area in bouton classes based on actin content relative to base: less than 25%, [-25%;0% [, [0%; +25% [, more than 25%. We did not observe any trend in bouton area in these bouton classes. However, we found that bouton area only was higher when actin in boutons is increased up to 25% when compared to base. Line represents the median. All data points are represented. **b-e**, n=14 larvae, 53 boutons. **f,** Schematics for quantifications of actin. We measured actin fluorescence intensity (integrated density) in bouton and at base at the initial frame (beginning) and at maximal size (when boutons stopped growing). We used as ROI a square 1x1 um. For more details see methods. Statistical significance was determined with non-parametric  Mann-Whitney test (two-tailed);*p<0.05, ns is non-significant.

### Supplementary Fig. 3

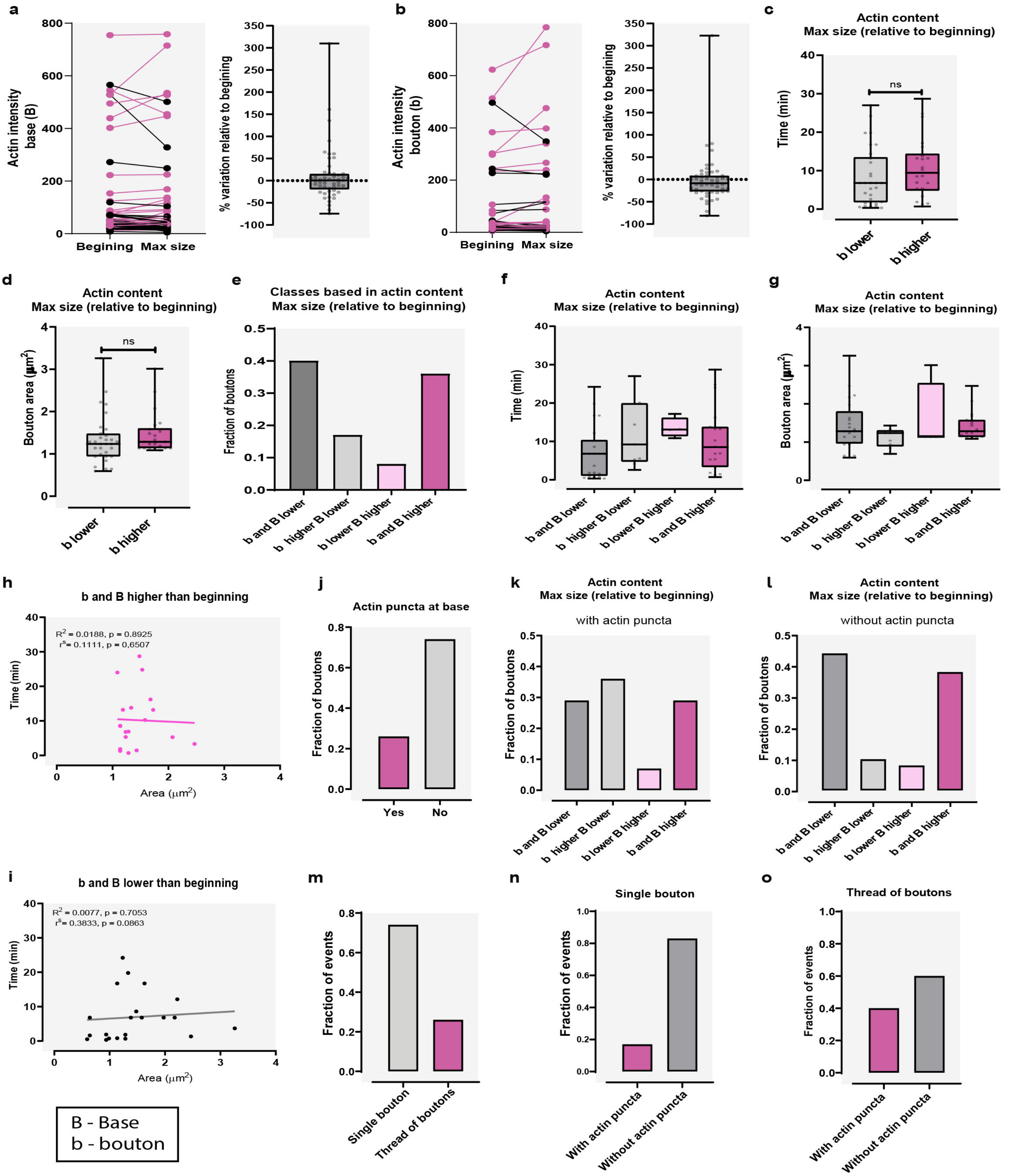


**Supplementary Fig. 3 Bouton classes based on actin changes in bouton and base and bouton formation dynamics. a, b,** Low variation in actin content in base (a) or in bouton (b) between beginning and max size. We show plots with absolute values for F-actin intensity (left) measured in beginning and correspondent max size, for bases (b) and boutons (B). Boxplot (min to max) showing % of variation (right) in F-actin intensity between beginning and max size (used as 100% reference). Line represents the median. All data points are represented. **c,** Boxplot (min to max) showing times of bouton formation was unchanged between boutons with actin lower or higher than beginning. Line is median. All data points are represented. **d,** Boxplot (min to max) showing bouton area was unchanged in boutons with actin lower or higher than beginning. Line is median. All data points are represented. **e,** Plot with fraction of boutons that at max size show (relative to beginning) actin: 1- lower in bouton and base; 2- lower in bouton and higher in base; 3- higher in bouton and lower in base; 4 – higher in bouton and in base. Boutons formed frequently from bases with low actin, acquiring less actin, or from bases with higher actin, acquiring more actin. **f,** Boxplot (min to max) showing times of bouton formation in classes 1-4. We did not observe any trend in bouton formation time in these classes. Line represents the median. All data points are represented. **g,** Boxplot (min to max) showing bouton area in classes 1-4. We did not observe any trend in bouton size in these classes. Line represents the median. All data points are represented. **h,i,** Plots with linear regression between time and size (area) for classes 1 (**i**) and 4 (**h**). We found no linear relationship between time and size for these classes. **j,** Plot with fraction of boutons where actin puncta was or not seen at base. **k,l,** Plot with fraction of boutons seen in classes 1-4 with (**k**) or without (**l**) actin puncta at the base. With actin puncta the proportion of boutons in class 2 was markedly increased. **m,** Plot with fraction of events observed that lead to formation of single bouton or threads of boutons. **n, o,** Plots with fraction of single bouton (**n**) or threads with boutons (**o**) seen with or without actin puncta. Actin puncta were more frequently observed in events resulting in the formation of threads of boutons. Data in this figure is representative of n= 14 larvae, 53 boutons. Statistical significance was determined with non-parametric Mann-Whitney test (two tailed); ns is non-significant.

#### Supplementary Fig. 4

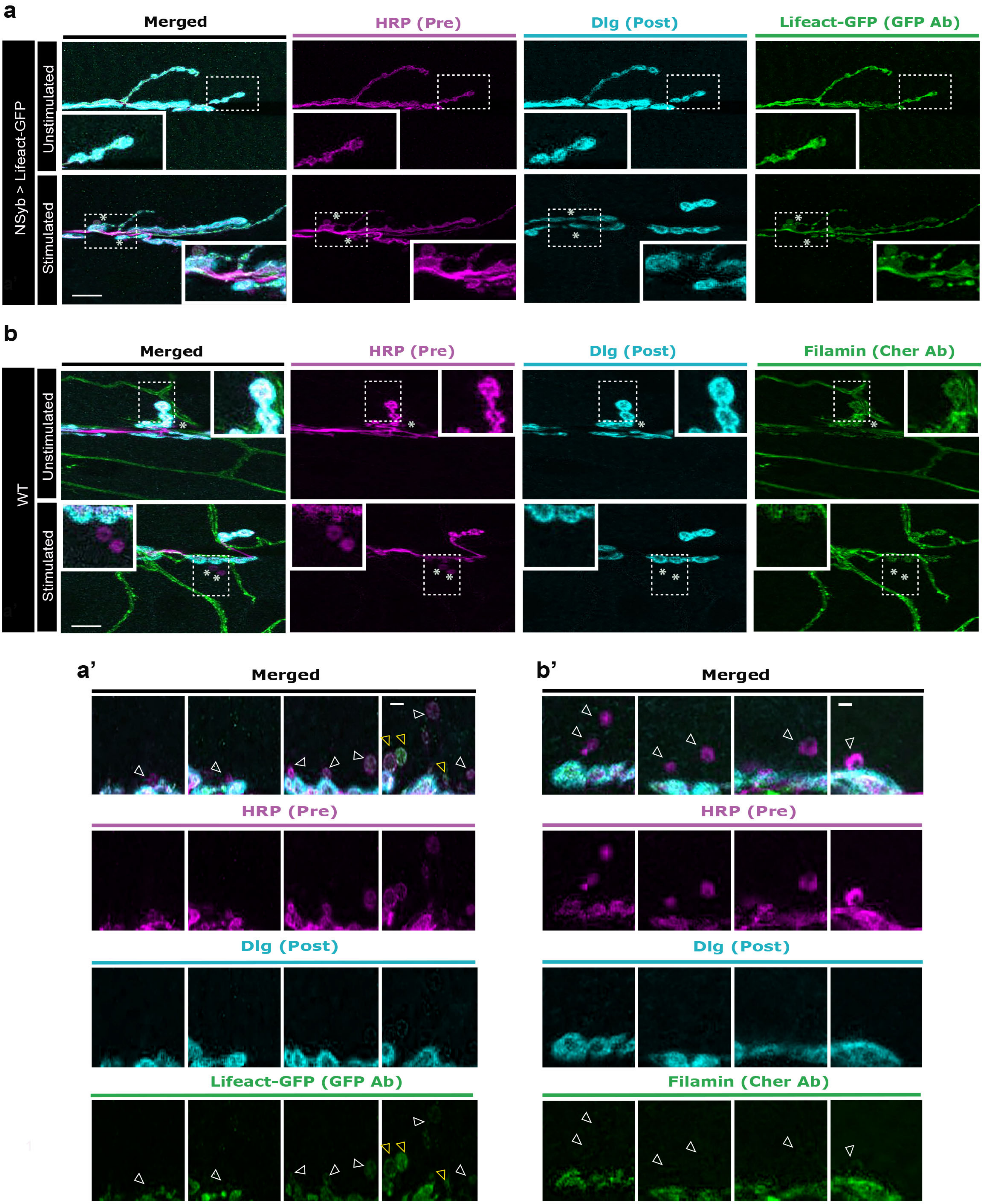


**Supplementary Fig. 4 F-actin and filamin localization at the NMJ before and after High-K+ stimulation. a,** We used animals expressing UAS-Lifeact-GFP under control of NSyb-Gal4 and did antibody staining for GFP to detect F-actin distribution in neurons. **a'** We could find new boutons without or with low amounts actin (white arrows) as well as actin-rich ones (yellow arrows), reminiscent of the blebs stages. **b,** To detect filamin at the WT NMJ we used an antibody directed against the presynaptic isoform of filamin. At the Drosophila NMJ the antibody immunoreactivity has been previously reported by GaYoung Lee & Tomas Schwartz (2016), with this isoform being present at the nerve, trachae, glia and puncta in presynaptic boutons. Our results reproduced this distribution (**b**). **b'** additionally, we saw that despite being present in MNs terminal boutons, filamin is not recruited to newly formed boutons (white arrows), suggesting that actin structure is weaker in the newly formed boutons. **a,b**, Unstimulated NMJ at the top. Stimulated NMJ on the bottom. Presynaptic membrane was labeled with HRP and postsynaptic membrane was labeled with Dlg. New boutons (asterisk) were identified by the lack of Dlg. Scale bar, 10 μM. Data in this figure is representative of n ≥ 6 larvae and ≥ 11 NMJs (unstim) or n ≥ 6 larvae and ≥ 15 NMJs (stim) and 2 biologically independent experiments for each staining.

#### Supplementary Fig. 5

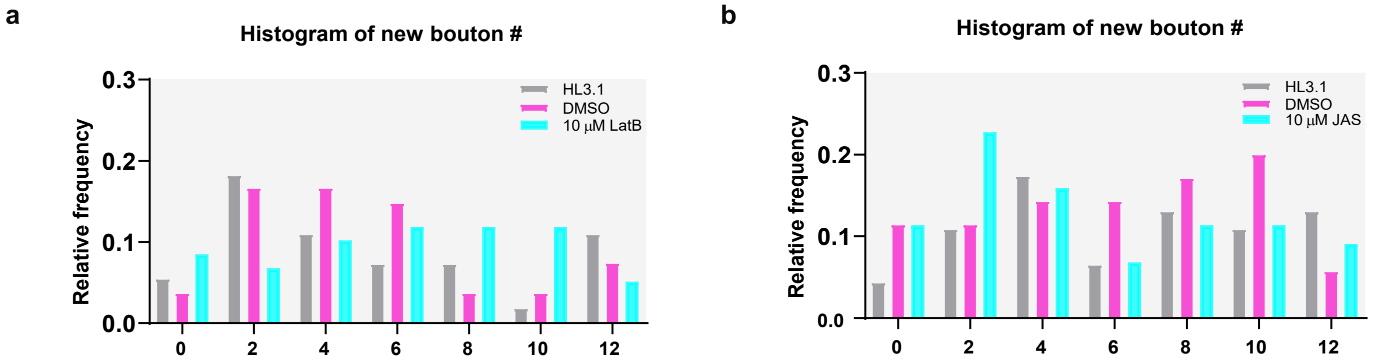


**Supplementary Fig. 5 New bouton number after manipulation of actin dynamics. a,** Histogram showing relative frequency new bouton number in WT NMJs treated with HL3.1, DMSO (solvent control ), and LatB (10 μM). n≥8 larvae, ≥50 NMJs for each condition and 3 biologically independent experiments. **b,** Histogram showing fraction of new boutons in WT NMJs treated with HL3.1, DMSO (solvent control), and JAS (10 μM). n≥6 larvae, ≥ 35 NMJs for each condition and 3 biologically independent experiments.

### Supplementary Fig. 6

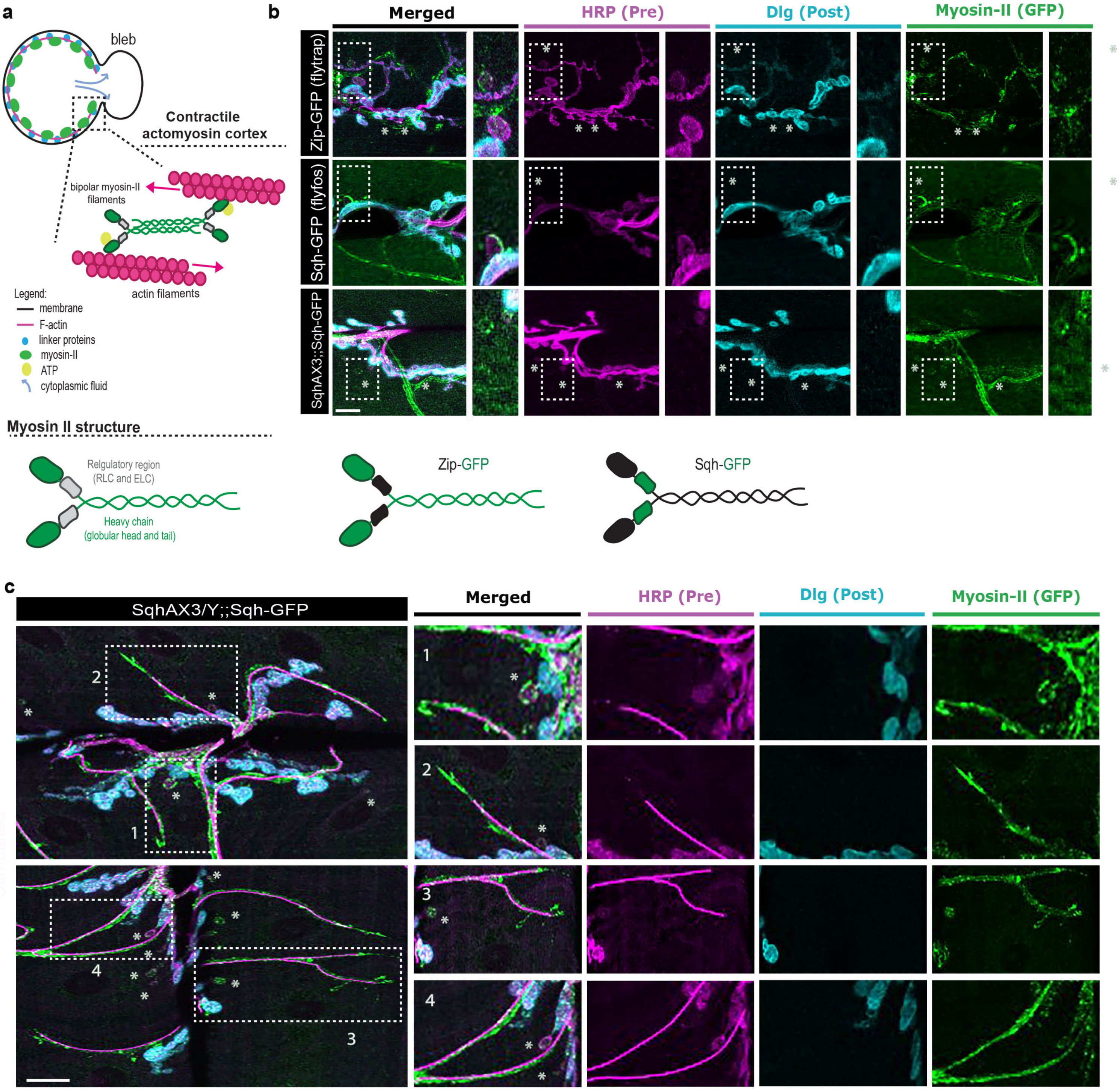


**Supplementary Fig. 6 Myosin-II-protein trap localization at the NMJ after High-K+ stimulation. a,** Schematics of cell's actomyosin cortex and myosin-II structure. Each molecule of myosin-II is composed of 2 heavy chains (Zip) and 2 regulatory light chains (Sqh). **b,** GFP protein traps of Sqh or Zip corroborate myosing cellular distribution observed with antibody staining **(Fig.3).** GFP-tagged Sqh in a Sqh-null background (SqhAX3) also showed normal myosin-II localization at the NMJ. Presynaptic membrane was labeled with HRP and postsynaptic membrane was labeled with Dlg. New boutons were identified by the lack of Dlg. Scale bar, 10 µM. **c,** NM-II localizes to trachea, as it co-localizes with autofluorescent (405 nm) tracheal lumen. Data in this figure is representative of 1 experiment for Zip-GFP and Sqh-GFP lines (at least 6 larvae, 10 NMJs per genotype) and at least 3 biologically independent experiments for SqhAX3;;Sqh-GFP (at least 6 larvae, ≥ 39 NMJs).

**Supplementary Fig. 7**

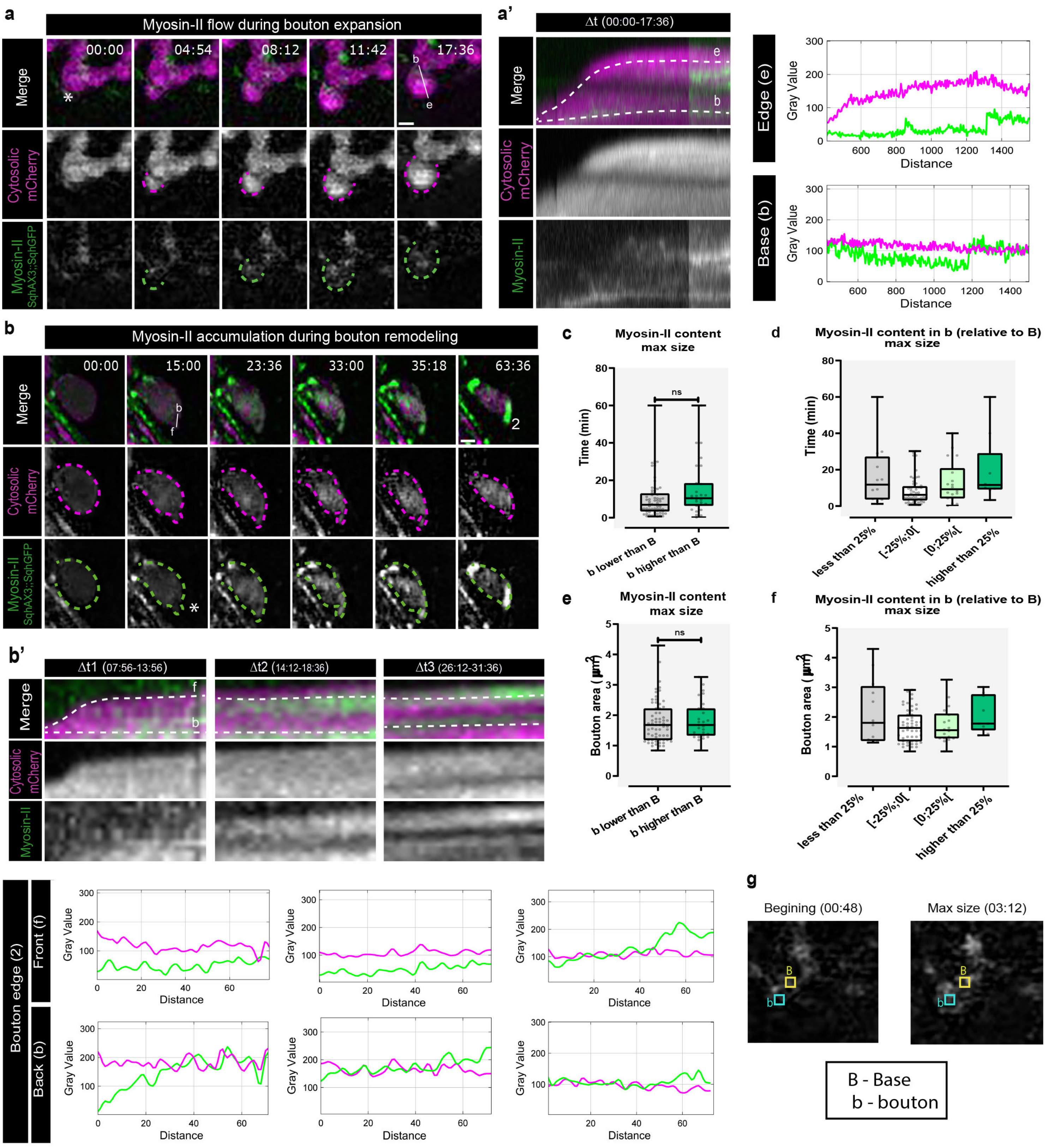


**Supplementary Fig. 7 Quantifications of myosin-II content in bouton and bouton formation dynamics. a, b** Time-lapse image of myosin-II dynamics during bouton formation. To follow bouton growth we used cytosolic mCherry under the control of LexA driver DvGlut and to label Myosin-II we used GFP-tagged Sqh (light chain) in a Sqh-null background (SqhAX3). Scale bar, 2 µm. Asterisk indicates where bouton will emerge. **a,** Example of myosin-II flow inside bouton during expansion. **b,** Example of myosin-II accumulation during bouton remodeling. **a', b'** Kymographs from **c** and **d.** We show kymographs for distinct Δt to highlight myosin-II dynamics. **a'** Shows that myosin-II is recruited to cortex of the growing bouton. We represent tracing lines along the bouton edge (e) and base (b) and the correspondent intensity values were plotted on the right. **b'** Small bouton forming at the edge (asterisk) of the parental bouton (Δt1) and that myosin-II clustered (represented as 2 on b) as the new bouton got smaller (Δt2 e Δt3). We represent dotted lines along the base back (b) and front (f) and the correspondent intensity values were plotted bellow. **c,** Boxplot (min to max) showing times of bouton formation were unchanged in boutons (b) with myosin-II lower or higher than base (B). Line is median. All data points are represented. **d,** Boxplot (min to max) showing times of bouton formation in bouton classes based on myosin-II content relative to base: less than 25%, [ -25%; 0[, [0; +25% [, more than 25%. Line represents the median. We noticed a non-statistical trend towards increased bouton formation time with higher myosin-II content. **e,** Boxplot (min to max) showing bouton area in boutons (b) with myosin-II lower or higher than base (B). Line is median. All data points are represented. **f,** Boxplot (min to max) showing bouton area in bouton classes based on actin content relative to base: less than 25%, [-25%;0% [, [0%; +25% [, more than 25%. We did not observe any trend in bouton area in these bouton classes. Line represents the median. All data points are represented. **c-f**, n=9 larvae, 92 boutons. **g**, Schematics for quantifications of myosin-II. We measured myosin-II fluorescence intensity (integrated density) in bouton and at base at the initial frame (beginning) and at maximal size (when boutons stopped growing). We used as ROI a square 1x1 um. Statistical significance was determined with non-parametric Mann-Whitney test (two-tailed); ns is non-significant.

### Supplementary Fig.8

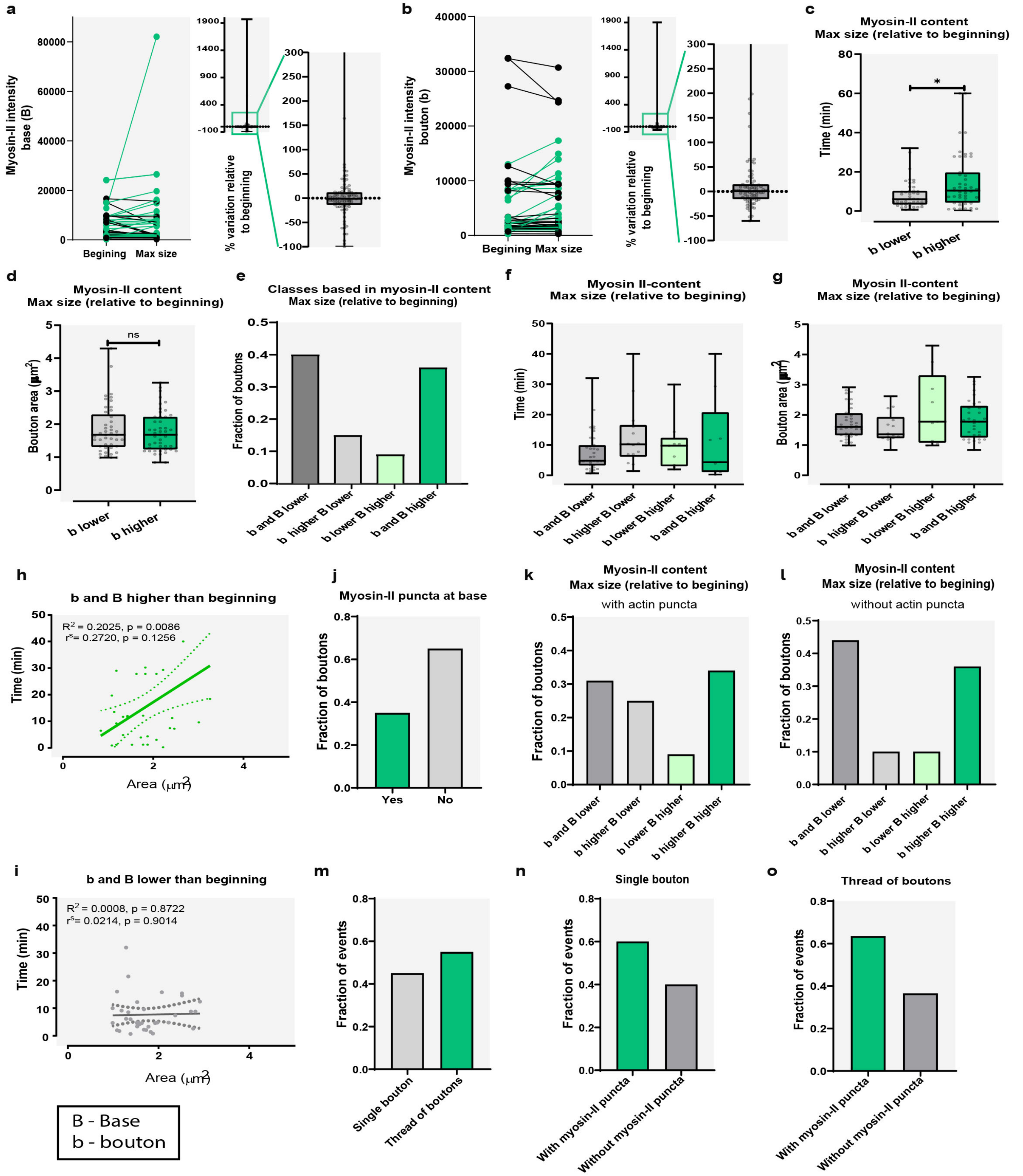


**Supplementary Fig. 8 Bouton classes based on myosin-II changes in bouton and base and bouton formation dynamics. a, b,** High variation in myosin-II content in base (a) or in bouton (b) between beginning and max size. We show plots with absolute values for myosin-II intensity (left) measured in beginning and correspondent max size, for bases (b) and boutons (B). We also present a boxplot (min to max) showing % of variation (right) in myosin-II intensity between beginning and max size (used as 100% reference). Line represents the median. All data points are represented. **c,** Boxplot (min to max) showing times of bouton formation was reduced in boutons with myosin-II lower than beginning, compared to boutons in which myosin-II was higher than beginning. Line is median. All data points are represented. **d,** Boxplot (min to max) showing bouton area was unchanged in boutons with actin lower or higher than beginning. Line is median. All data points are represented. **e,** Plot with fraction of boutons that at max size show (relative to beginning) actin: 1- lower in bouton and base; 2- lower in bouton and higher in base; 3- higher in bouton and lower in base; 4 – higher in bouton and in base. Boutons formed frequently from bases with low myosin-II, acquiring less myosin-II, or form bases with higher myosin-II, acquiring more myosin-II. **f,** Boxplot (min to max) showing times of bouton formation in classes 1-4. We did not observe any trend in bouton formation time in these classes. We noticed a non-statistical trend towards increased bouton formation time with higher myosin-II content Line represents the median. All data points are represented. **g,** Boxplot (min to max) showing bouton area in classes 1-4. We did not observe any trend in bouton size in these classes. Line represents the median. All data points are represented. **h,i,** Plots with linear regression between time and size (area) for classes 1 (**i**) and 4 (**h**). We find a weak linear relationship between time and size for class 4; no association was found for class 1. **j,** Plot with fraction of boutons where myosin-II puncta was or not seen at base. **k,l,** Plot with fraction of boutons seen in classes 1-4 with (**k**) or without (**l**) actin puncta at the base. With actin puncta the proportion of boutons in class 2 was markedly increased. **m,** Plot with fraction of events observed that lead to formation of single bouton or threads of boutons. **n, o,** Plots with fraction of single bouton (n) or threads with boutons (o) seen with or without myosin-II puncta. We found no differences in events resulting in the formation of threads of boutons when myosin-II puncta was observed at base. Data in this figure is representative of n= 9 larvae, 92 boutons. Statistical significance was determined with non-parametric two-tailed Mann-Whitney test; *p<0.05, ns non-significant.

#### Supplementary Fig. 9

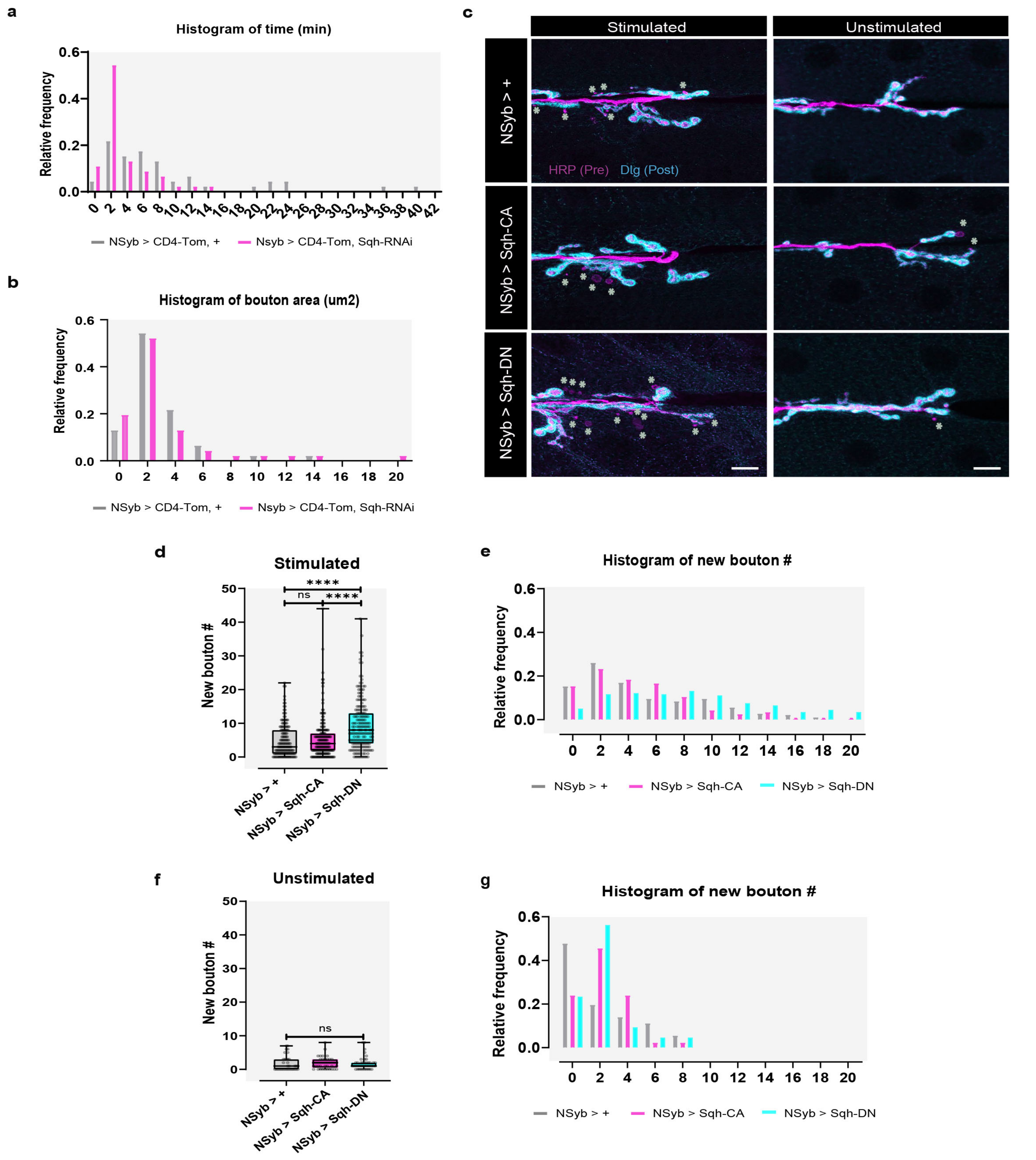


**Supplementary Fig. 9 Effects of reducing myosin-II or altering activity of myosin-II in activity-dependent plasticity. a,b** Histograms of relative frequency for times of bouton formation (a) and bouton area (b) in NSyb-Gal4,CD4-Tom/+ and in NSyb-Gal4,CD4-Tom/Sqh-RNAi. n = 5 larvae, 46 boutons (for Sqh-RNAi) and 13 larvae, 46 boutons (for control). **c**, Images of control animals and animals expressing constitutively active (CA) or dominant negative (DN) forms of myosin-II with (left) or without (right) High-K+ stimulation. We used NSyb-Gal4 (pan-neuronal driver) to express in MNs Sqh-CA (phosphomimic) or Sqh-DN (non phosphorylatable). Presynaptic membrane was labeled with HRP and postsynaptic membrane was labeled with Dlg. New boutons (asterisk) were identified by the lack of Dlg. Scale bar, 10 µm. **d,** Boxplot (min to max) showing that neuronal expression of Sqh-DN, but not Sqh-CA, increased new bouton frequency after acute stimulation, compared to control. Line is at median. All data points are represented. **e,** Histogram showing relative frequency of new boutons in stimulated NMJs. **d,e** n ≥ 30 larvae, ≥ 197 NMJs for each line and at least 5 biologically independent experiments. **f,** Boxplot (min to max) showing that neuronal expression of Sqh-CA or Sqh-DN did not change new bouton frequency during development (unstimulated control). Line is at median. All data points are represented. **g,** Histogram showing that frequency of new boutons in unstimulated NMJs. **f, g** n ≥6 larvae, ≥ 35 NMJs for each line and 3 biologically independent experiments. Statistical significance was determined with non-parametric Kruskal-Wallis test (two-tailed);****p<0.0001, ns is non-significant. Significant changes observed in the distribution of new bouton number were seen across all experiments, rather in every single experiment.

#### Supplementary Fig.10

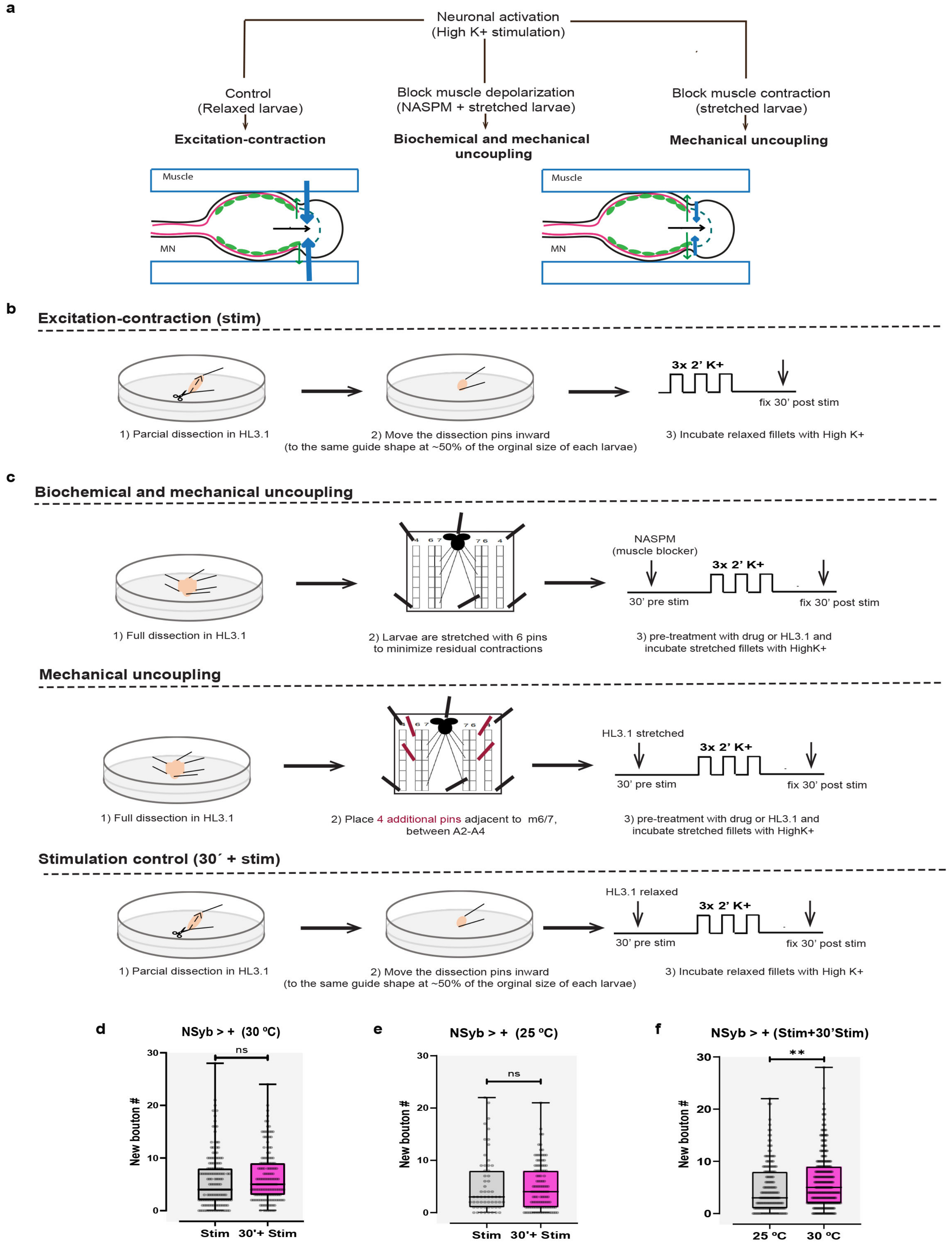


**Supplementary Fig. 10 Experimental design for blocking muscle contractions during activity-dependent plasticity and controls. a,** Schematics showing strategies adopted to uncouple MN excitability from muscle contraction: muscle blocker vs mechanically stretching the larvae. **b,** Schematics for normal stimulation protocol (stim). **c,** Schematics for protocols for stimulations uncoupling excitation-contraction and stimulation control that was done in parallel for each experiment (30'+ stim). **d**, New bouton number was unchanged between stim and 30'+ stim for control (NSyb > +) animals raised at 30ºC. **e**, New bouton number was unchanged between stim and 30'+ stim for control animals raised at 25ºC. **f**, New bouton number was increased in control animals raised at 30ºC, compared with the ones that stayed at 25ºC. Line is at median. All data points are represented. n ≥25 larvae, ≥ 159 NMJs (30ºC) or n ≥ 10 larvae, ≥ 59 NMJs (25ºC) and at least 5 biologically independent experiments. Statistical significance was determined with non-parametric Kruskal--Wallis test (two-tailed);****p<0.0001, ns is non-significant.

#### Supplementary Fig. 11

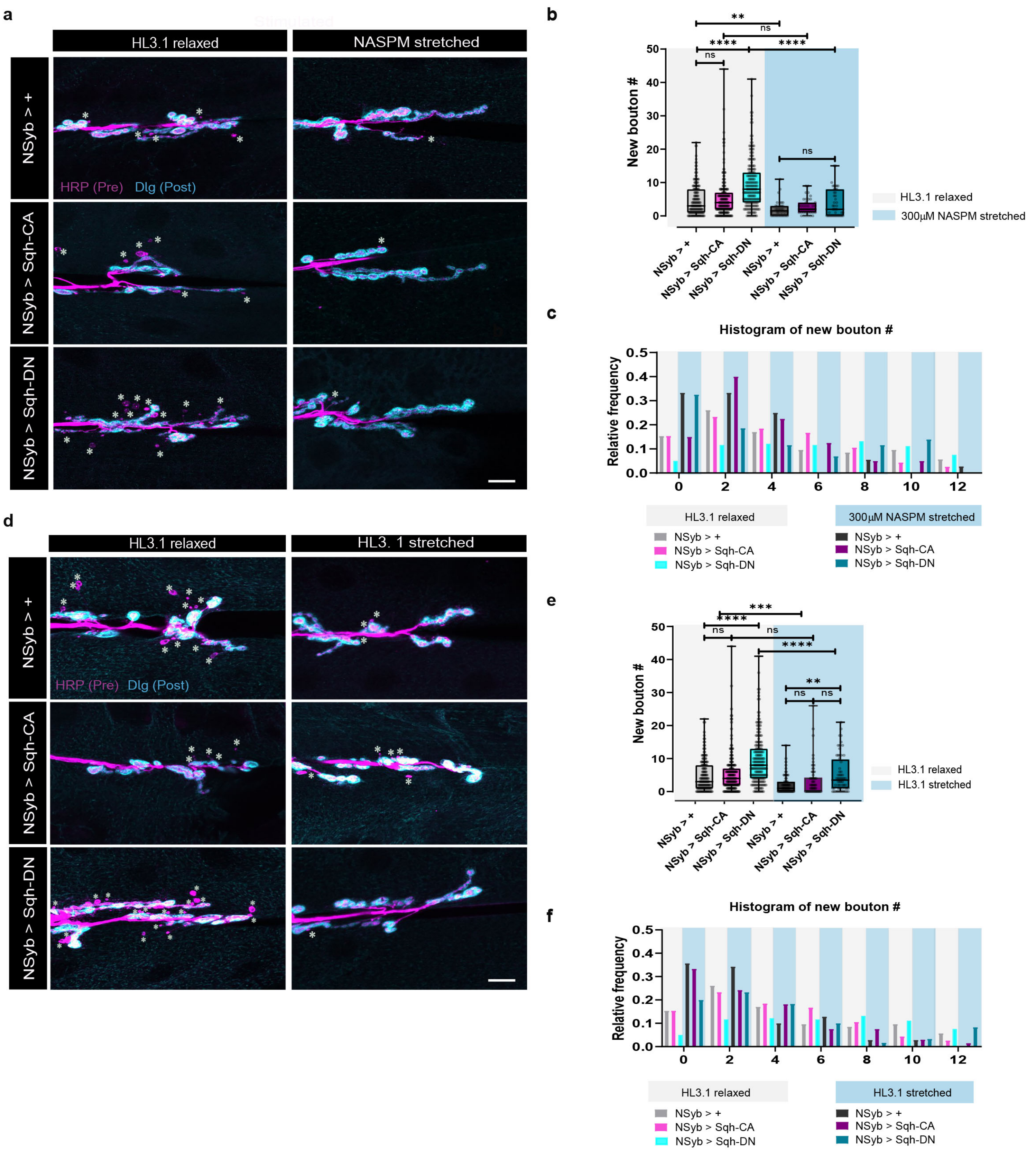


**Supplementary Fig. 11 Effects of blocking muscle activation and/or contraction in activity-dependent bouton formation with Myosin-II activation or inactivation. a,b,** Blocking muscle contraction with NASPM decreased activity-dependent formation in control and in neuronal Sqh-DN, but not in Shq-CA. **b,** Boxplot (min to max) of new bouton number. Line is at median. All data points are represented. **c,** Histogram relative frequency of new bouton numbers for animals treated with HL3.1 or NASPM. n≥6 larvae, ≥ 36 NMJs and 5 biologically independent experiments. **d,e,** Blocking muscle contraction by mechanically stretching the larvae decreased activity-dependent formation in control and in neuronal Sqh-DN, but not in Shq-CA. **e,** Boxplot (min to max) of new bouton number. Line is at median. All data points are represented. **f,** Histogram relative frequency of new bouton numbers for relaxed or stretched animals. n≥15 larvae, ≥ 60 NMJs and 5 biologically independent experiments. Presynaptic membrane was labeled with HRP and postsynaptic membrane was labeled with Dlg. New boutons (asterisk) were identified by the lack of Dlg. Scale bar, 10 µM. Statistical significance was determined with non-parametric Kruskal-Wallis test (two-tailed);****p<0.0001,***p<0.001, **p<0.01, ns is non-significant.
